# HIFα regulates developmental myelination independent of autocrine Wnt signaling

**DOI:** 10.1101/2020.03.30.015131

**Authors:** Sheng Zhang, Yan Wang, Jie Xu, Wenbin Deng, Fuzheng Guo

**Author notes:** **Corresponding author:** Fuzheng Guo, PhD, 2425 Stockton Blvd, Sacramento, CA 95817.

## Abstract

The developing CNS is exposed to physiological hypoxia, under which hypoxia inducible factor alpha (HIFα) is stabilized and plays a crucial role in regulating neural development. The cellular and molecular mechanisms of HIFα in developmental myelination remain incompletely understood. Previous concept proposes that HIFα regulates CNS developmental myelination by activating the autocrine Wnt/β-catenin signaling in oligodendrocyte progenitor cells (OPCs). Here, by analyzing a battery of genetic mice of both sexes, we presented *in vivo* evidences supporting an alternative understanding of oligodendroglial HIFα-regulated developmental myelination. At the cellular level, we found that HIFα was required for developmental myelination by transiently controlling upstream OPC differentiation but not downstream oligodendrocyte maturation and that HIFα dysregulation in OPCs but not oligodendrocytes disturbed normal developmental myelination. We demonstrated that HIFα played a minor, if any, role in regulating canonical Wnt signaling in the oligodendroglial lineage or in the CNS. At the molecular level, blocking autocrine Wnt signaling did not affect HIFα-regulated OPC differentiation and myelination. We further identified HIFα-Sox9 regulatory axis as an underlying molecular mechanism in HIFα-regulated OPC differentiation. Our findings support a concept shift in our mechanistic understanding of HIFα-regulated CNS myelination from the previous Wnt-dependent view to a Wnt-independent one and unveil a previously unappreciated HIFα-Sox9 pathway in regulating OPC differentiation.

## Introduction

HIFα is a master transcriptional regulator of the adaptive response to hypoxia (Semenza, 2012). The oxygen concentration in the developing CNS was reported ranging from 0.5 to 7% (Ivanovic, 2009; Zhang et al., 2011). HIFα protein (HIF1α and HIF2α) is constitutively translated but subjected to rapid turnover (half-life<5 min) by the proteasome-mediated degradation in which the von Hippel-Lindau (VHL) protein plays an essential role (Semenza, 2012). Under physiological or pathological hypoxia or VHL mutation, HIFα accumulates in the nucleus where it complexes with the stable HIFβ subunit and other co-activators to activate target gene expression.

The role of HIFα, particularly the representative HIF1α, in neural precursor cells under physiological conditions has been extensively investigated (Cunningham et al., 2012; Li et al., 2014; Milosevic et al., 2007; Tomita et al., 2003). Aggravated hypoxia and/or ischemia causes severe disturbance of normal oligodendroglial myelination in the preterm infants born between 28^th^ and 37^th^ week gestational ages (Volpe, 2009). Animal studies demonstrate that pathological hypoxia insult delays developmental myelination in preterm-equivalent early postnatal murine CNS (Jablonska et al., 2012; Liu et al., 2011; Liu et al., 2013). The role of HIFα in developmental myelination has not been defined until an important study reported that HIFα plays a major role in hypoxia-elicited myelination disturbance in cell/brain slice culture systems (Yuen et al., 2014). However, the cellular and molecular mechanisms underlying oligodendroglial HIFα-regulated myelination remain incompletely defined. The current popular hypothesis states that HIFα dysregulation disturbs developmental myelination by activating autocrine Wnt/β-catenin signaling (Yuen et al., 2014). This “Wnt-dependent” mechanistic hypothesis is in line with the inhibitory effect of intracellular Wnt/β-catenin activation on CNS myelination (Guo et al., 2015), but it was only tested in the cell / slice culture systems in combination with pharmacological manipulations. Given the intrinsic caveats of non-cellular specificity and/or off-target effects of pharmacological application, it is imperative to determine, by using *in vivo* genetic approaches, the involvement of oligodendroglial HIFα-derived Wnt signaling in developmental myelination.

CNS developmental myelination consists of at least two major sequential steps: OPC differentiation into oligodendrocytes (OLs, referred to as OPC differentiation) and oligodendrocyte maturation and myelination (Guo et al., 2015; Huang et al., 2013). In humans, developmental myelination starts in the third trimester of gestational ages, which is equivalent to the perinatal and early postnatal ages in rodents (Semple et al., 2013). Therefore, we used the murine CNS of perinatal and early postnatal ages as a third trimester-equivalent *in vivo* model to dissect the cellular and molecular mechanisms underlying development myelination. By analyzing a series of cell-specific HIFα and Wnt genetic mutant mice, we provided convincing evidence supporting an alternative HIFα model of CNS myelination: HIFα transiently regulates developmental myelination by controlling OPC differentiation but subsequent OL maturation and its hyperactivation disturbs OPC differentiation in a manner independent of autocrine Wnt/β-catenin signaling. Our results further demonstrate that sustained Sox9 activation is a downstream mechanism underlying disturbed OPC differentiation elicited by HIFα hyperactivation.

## Materials and Methods

### Transgenic animals

The following transgenic mice were used in our study: *Cnp-Cre* mice (RRID: MGI_3051754) kindly provided by Dr. Nave (Lappe-Siefke et al., 2003), *Sox10-Cre* (RRID: IMSR_JAX:025807), *Sox10-CreER^T2^* (RRID:IMSR_JAX: 027651), *Pdgfrα-CreER^T2^* (RRID: IMSR_JAX:018280), Hif1α-floxed (RRID: IMSR_JAX:007561), Hif2α-floxed (RRID: IMSR_JAX:008407), *Vhl-floxed* (RRID: IMSR_JAX:012933), and Wls-floxed (RRID: IMSR_JAX:012888). In our study, we found no difference in oligodendroglial phenotypes between non-Cre mice carrying floxed alleles and Cre transgenic mice carrying no floxed alleles. Therefore, we used non-Cre and/or Cre transgenic mice from same litters of cKO as littermate control mice. All mice were maintained on the C57BL/6 background and housed in 12 h light/dark cycle with free access to water and food. Animal protocols were approved by the Institutional Animal Care and Use Committee at the University of California, Davis.

### Tamoxifen treatment

Tamoxifen (TM) (T5648, Sigma) was prepared at a concentration of 30 mg/ml in a mixture of ethanol and sunflower seed oil (1:9, v/v). All study mice including littermate control and conditional knockout were received tamoxifen intraperitoneally at a dose of 200 μg/g body weight.

### Hypoxia/ischemia (H/I)-induced brain white matter injury

We used our established protocols (Chung et al., 2013; Shen et al., 2010) to induce H/I injury in C57BL/6J background mice at postnatal day 6 (P6), a time point when OPC differentiation barely occurs in the brain. In brief, P6 mice were anesthetized with indirect cooling by wet ice, which were immediately subjected to permanent occlusion (by cauterizers) of the right side common carotid artery (ischemia). One hour after the artery occlusion, mice were exposed to 6% O_2_ for 45min (hypoxia). After hypoxia exposure, the mice were returned to the dam. Sham operation was performed in the same way as H/I injury including anesthesia, neck skin incision, and exposure of the right common carotid artery but without artery cauterization or hypoxia. Previous study demonstrates that H/I injury results in widespread periventricular white matter injury including arrested OPC differentiation and microglial/astroglial activation (Liu et al., 2011).

### Tissue processing and immunohistochemistry

After anesthetization by ketamine/xylazine mixture, mice were perfused with cold PBS. Tissues were collected, fixed with fresh 4% PFA for 2 h at room temperature, cryopreserved in 30% sucrose overnight at 4°C and embedded in OCT (VWR International, Wetzlar, Germany). Serial coronal sections (14 μm) were cut by a Leica Cryostat (CM 1900-3-1). Tissue sections were incubated with primary antibody overnight at 4°C after blocking with 10% donkey serum at room temperature for at least 1 h. After 3 times wash, the fluorescence conjugated secondary antibody (Jackson ImmuneResearch) was applied and incubated for 1.5 h. Then the slice was protected by mounting medium after DAPI nucleus staining. The following primary antibodies were used in immunohistochemical study: HIF-1α (RRID:AB_409037,1:200,Cayman Chemical #100642,), Sox10 (RRID: AB_2195374,1:100,Santa Cruz Biotechnology, sc-17342; RRID: AB_778021, 1:500, Abcam, ab27655), Olig2 (RRID: AB_570666,1:200, Millipore, AB9610), Mbp (RRID:AB_2564741, 1:500, Biolegend, SMI-99,), SMI312 (RRID:AB_2135329, 1:1000, Covance, SMI-312R), CC1 (RRID:AB_213434, 1:200, Calbiochem, OP80,), PDGFRa (RRID:AB_2236897,1:200, R&D system, AF1062), SMI32 (RRID:AB_2314912, 1:1000, Biolegend, SMI-32P), and Sox9 (RRID:AB_2194160,1:200, R&D system, AF3075).

### In situ hybridization and transmission electron microscopy (TEM)

Briefly, the slices were treated with Proteinase K (Thermo Fisher, AM2548) and acetylated by triethanolamine (Sigma, 90279) and acetic anhydride (Sigma, 320102). The tissue was incubated with 100 ng/ul DIG labelled c-RNA at 65^0^C overnight. Then the slices were incubated with Alkaline phosphatase (AP) anti-DIG secondary antibody (1:100,11093274910, Sigma) overnight after blocking with 10% donkey serum. The signal was determined by nitroblue tetrazolium (NBT) and 5-bromo-4-chloro-3-indolyl phosphate (toluidine salt) (BCIP) (Sigma, 72091).Semithin sections for toluidine blue myelin staining and ultrathin section for TEM imaging were prepared according to our previous protocols (Zhang et al., 2018a; Zhang et al., 2018b). TEM imaging was conduction by the professionals at UC Davis Core Facility who were blinded to the mouse genotypes.

### Enzyme-linked immunosorbent assay (ELISA)

The Wnt3a protein level was measured by the mouse Wnt3a ELISA kit (M13146365, CSB-EL026136MO, Cusabio) according to the manufacturer’s instruction.

### Primary OPC culture and gene manipulation

The mouse cortex between P0 and P2 was collected and dissociated with papain kit and DNase I (#LK003176, Worthington; 250U/ml; #D5025, Sigma). OPCs were purified by immnopanning approach according to our previous protocol (Lang et al., 2013). Purified OPCs were used for experiments of Wls knockdown, Wnt3a overexpression, and Sox9 overexpression. Wls-shRNA (TRCN 0000 234932, Mission ShRNA bacterial Glycerol stock NM_026582) and scramble control (Mission TRC2 PlkO.5-PURO Non-Mammalian shRNA control Plasmid) were purchased from Sigma-Aldrich. Wnt3a plasmid pLCN-Wnt3a-HA (Addgene, #18030) and empty pLCN-exp (Addgene, #64865) were used for Wnt3a overexpression experiment. The transfection of Wls-shRNA and pLCN-Wnt3a-HA and controls was performed by using HiPerFect transfection reagent (#301704, Qiagen) according to the manual. p-WPXL-Sox9 (Addgene, #36979) and control plasmid p-WPXL (Addgene, #12257) were used for Sox9 overexpression and the transfection was performed by using FuGENE6 Transfection reagent (Promega, #E2691, lot#000371257).

Primary rat OPCs were used in Sox9 knockdown and DMOG treatment. DMOG (0.5 mM dissolved in DMSO, D3695, Sigma), and Sox9 siRNA (SASI_Rn02_00372739, Sigma) or negative control siRNA (SIC001-5X1NMOL, Sigma) were applied according to the experimental design in Figure 12H. Transfection of Sox9 siRNA was conducted by using HiPerFect transfection reagent (#301704, Qiagen).

### Purification of brain OPCs by magnetic activated cell sorting (MACS)

MACS purification of OPCs was performed according to the protocols in the kit instruction from Miltenyi Biotec. The mouse forebrain was dissociated using the papain dissociation system (LK003176, Worthington) and gentleMACS™ Dissociator (Miltenyi Biotec, #130-092-235). Astrocytes and microglia in the cell suspension were removed by using anti-ACSA-2 micro beads (130-097-679, Miltenyi Biotec) and anti-CD11b micro beads (130-049-601, Miltenyi Biotec), respectively. After astroglial and microglial removal, the cell suspension was then incubated with anti-O4 micro beads (130-094-543, Miltenyi Biotec) for OPC purification. O4^+^ cells were collected for RNA preparation.

### RNA preparation, reverse transcription and real-time quantitative PCR (RT-qPCR)

Total RNA was obtained using Qiagen RNeasy for lipid tissue kit (74804, Qiagen) according to the manual. We used on-column DNase I digestion to eliminate DNA contamination. Complementary DNA (cDNA) was prepared using Qiagen Omniscript RT Kit (205111, Qiagen). Mouse and rat gene expression was normalized to the internal control Hsp90 and β-actin, respectively, and calculated by the equation of 2^^(Ct(cycle threshold) of Hsp90 - Ct of indicated genes)^. The value of control groups was normalized to 1 throughout the study. Real time qPCR was done by using QuantiTect SYBR® Green PCR Kit (204145, QIAGEN) and Stratagene MP3005P thermocycler. The qPCR primers used in the study were shown as below.

Mouse Wls: AGGGCAAGGAAGAAGGAGAG / ATCCCTCCAACAATGCAGAG
Mouse Glut1: CAGTTCGGCTATAACACTGGTG / GCCCCCGACAGAGAAGATG
Mouse Ldha: CATTGTCAAGTACAGTCCACACT / TTCCAATTACTCGGTTTTTGGGA
Mouse Hk2: TGATCGCCTGCTTATTCACGG / AACCGCCTAGAAATCTCCAGA
Mouse Axin2: AACCTATGCCCGTTTCCTCTA / GAGTGTAAAGACTTGGTCCACC
Mouse Naked1: CAGCTTGCTGCATACCATCTAT / GTTGAAAAGGACGCTCCTCTTA
Mouse Wnt7a: CGACTGTGGCTGCGACAAG / CTTCATGTTCTCCTCCAGGATCTTC
Mouse Wnt7b: CTTCACCTATGCCATCACGG / TGGTTGTAGTAGCCTTGCTTCT
Mouse Vegfa: GCACATAGAGAGAATGAGCTTCC / CTCCGCTCTGAACAAGGCT
Mouse Mbp: ACACGAGAACTACCCATTATGGC / CCAGCTAAATCTGCTGAGGGA
Mouse Plp (exon3b-specific): CCAGAATGTATGGTGTTCTCCC / GGCCCATGAGTTTAAGGACG
Mouse Cnp: TTTACCCGCAAAAGCCACACA / CACCGTGTCCTCATCTTGAAG
Mouse Mobp: AACTCCAAGCGTGAGATCGT / CAGAGGCTGTCCATTCACAA
Mouse Opalin: CTGCCTCTCACTCAACATCA / GCTGGATCAAAGTAAACAGC
Mouse Mog: AGCTGCTTCCTCTCCCTTCTC / ACTAAAGCCCGGATGGGATAC
Mosue Qk: CTGGACGAAGAAATTAGCAGAGT / ACTGCCATTTAACGTGTCATTGT
Mouse Myrf: CAGACCCAGGTGCTACAC / TCCTGCTTGATCATTCCGTTC
Mouse Sox9: AGTACCCGCATCTGCACAAC / ACGAAGGGTCTCTTCTCGCT
Mouse Hsp90: AAACAAGGAGATTTTCCTCCGC / CCGTCAGGCTCTCATATCGAAT
Rat Bnip3: GCTCCCAGACACCACAAGAT / TGAGAGTAGCTGTGCGCTTC
Rat Mbp: TTGACTCCATCGGGCGCTTCTTTA / TTCATCTTGGGTCCTCTGCGACT
Rat Sox9: CTGAAGGGCTACGACTGGAC / TACTGGTCTGCCAGCTTCCT
Rat β-actin: CGTCTTCCCCTCCATCGT / GGAGTCCTTCTGACCCATACC

### Chromatin immunoprecipitation (ChIP) by HIFα antibody and qPCR verification

ChIP was performed using the SimpleChIP plus Enzymatic Chromatin IP kit (Cell Signaling Technology, #9005) following the manufacturer’s protocol with some modifications. Briefly, primary cultured rat OPCs (4 x 10^6^ cells) treated with 1 mM DMOG for 7 hours were crosslinked in culture medium containing 1% formaldehyde at room temperature for 10 min. After addition of glycine solution to stop the reaction, the cells were then collected, centrifuged and lysed. Nuclei were collected and treated with 0.5 ul micrococcal nuclease per IP for 20 min at 37 °C. The reaction was stopped using 0.05 M EDTA, and nuclear membranes were destroyed by sonication. Then the supernatant was collected by centrifugation. Chromatin solutions were incubated with 10 ug of anti-HIF1α (Cayman, Rabbit, #10006421) antibody or anti-rabbit IgG (Cell Signaling #2729) overnight at 4 °C, and immunoprecipitated with protein G magnetic beads for 2 hours at 4 °C with rotation. Bound DNA-protein complexes were washed and eluted, and crosslinks were reversed according to kit instructions. DNA was purified using spin columns and used for ChIP-qPCR analysis. The fold enrichment of specific genomic regions was assessed relative to the qPCR data from the IPs normalized to the control IgG values. Fold enrichment = 2^Λ-[(Ct IP) -(Ct IgG)]^. qPCR primer sets for ChIP binding site verification are:

Negative site: forward TTCCTCTTGGGATGGTTGTC / reverse CCACCTCTGGGGAGTATGAA
−75bp site: forward TTTCAAAATCCGGTCCAATC / reverse CCCCCTTCACCTTAGAGACC
−828bp site: forward CGGGGAAGGACTTGTCAGT / reverse ATGAAAACCAAAGCCAAGCA
−1211bp site: forward CGGCTCCAAGCACTCTTAAA /reverse GCGTCCTTTTAGACCTGCAC
−1928 site: forward GTCGTTCTCGCTGCCTTTAG / reverse TTGAAGAGACCAGGGACCAC
−2098 site: forward CT CTGGAT GTT GCCGAAAAT / reverse CTCCACGCAAGCGTTTTTAT

### Western blot and antibodies

Twenty microgram protein was loaded into AnykD Mini-PROTEAN gel (#4569035, BIO-RAD). After blocking, the membrane was incubated with primary antibodies overnight. The membrane was washed 3 times by TBST (TBS with 1% tween 20). Then the HRP-conjugated secondary antibody was reacted with membrane for 1.5 hours. Protein signals were developed by using ECL kit (NEL103001EA, Perkin Elmer). Quantification of interested protein bands were conducted by using NIH Image J software. Antibodies used in western blot are MBP (SMI-99, RRID:AB_2564741, 1:2000, Biolegend), Active β-catenin (#05­665, RRID: AB_309887, 1:1000, Millipore), Axin-2(#6163, RRID: AB_10904353, 1:1000, Prosci) and β-actin (3700, RRID:AB_2242334, 1:2000, Cell signaling Technology).

### Experimental design and statistical analyses

Data collection was conducted by lab members who were blinded to mouse genotypes. We used Shapiro-Wilk approach to test data normality. F test and Browne-Forsythe test were used to variance equality of two groups and three or more groups, respectively. We used scatter graphs to present the quantification data throughout the current manuscript. Both male and female mice were included in our data analysis. All data were presented as mean ± standard effort of mean (s.e.m.) where each dot represents one mouse. Unpaired two-tailed Student’s t test was used for statistically analyzing two groups of data. The *t*-values and the degree of freedom (df) were shown as t_(d_f) in each graph. One-way ANOVA followed by Tukey’s post-hoc test was used for statistically analyzing three or more groups of data. The F ratio, and DFn and DFd was presented as F_(D_Fn, DFd) in the figure legends where DFn stands for degree of freedom numerator and DFd for degree of freedom of denominator. The exact P-value of one-way ANOVA test was also described in the figure legends. All data graphing and statistical analyses were performed using GraphPad Prism version 8.0. The P-value was designated as * p < 0.05, ** *p* < 0.01, *** *p* < 0.001, ns, not significant p>0.05.

## Results

### Transient HIFα stabilization during early CNS development

HIF1α and HIF2α share high sequence homology and function in a similar manner through partnering with HIF1β and subsequently activating HIFα target genes (Semenza, 2012). Therefore, we used antibodies against the representative HIF1α to determine whether HIFα protein is stabilized in the perinatal and early postnatal CNS, where the vasculature is still in immature state (Harb et al., 2013; Yuen et al., 2014). We found that HIF1α protein was detected by immunohistochemistry (IHC) in Sox10^+^ oligodendroglial lineage cells in the white matter areas of the spinal cord (Fig. 1A) and the forebrain (Fig. 1C) at postnatal day 5 (P5). As positive controls, HIF1α immunoreactive signals were markedly increased in age-matched transgenic *Sox10-Cre:Vhl^fl/fl^* mice (Fig. 1B, D), in which Sox10-Cre-mediated VHL disruption prevents HIFα from degradation and causes HIFα accumulation specifically in oligodendroglial lineage cells.

**Figure 1.**
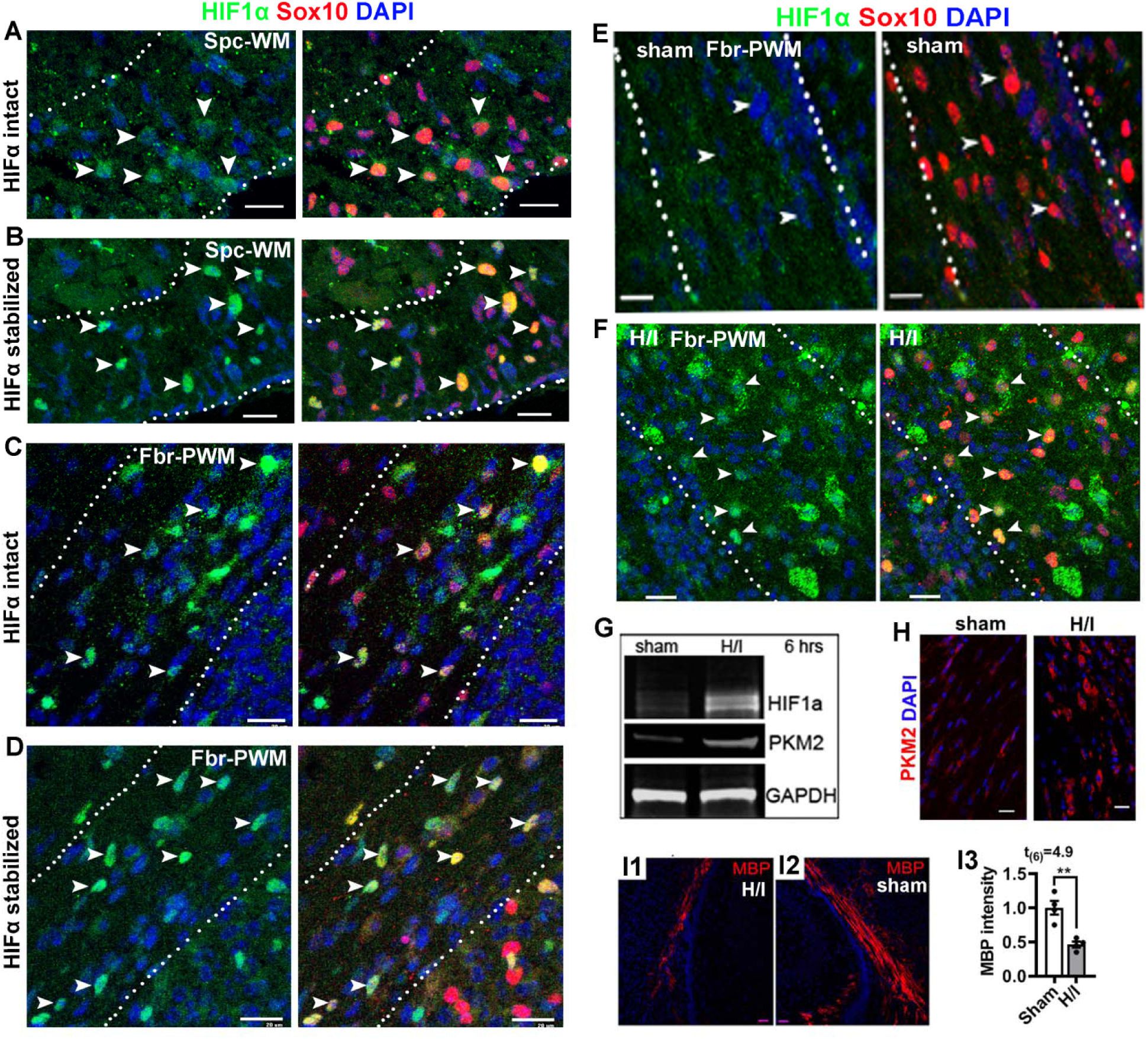
HIFα stabilization in the early postnatal murine CNS. **A-B**, immunohistochemistry (IHC) of HIF1α and oligodendroglial lineage marker Sox10 in the spinal cord ventral white matter (Spc-WM, marked by dotted lines) of HIFα intact mice **(A)** and HIFα stabilized *(Sox10-Cre:Vh*P^m^) mice **(B)** at postnatal day 5 (P5). **C-D**, IHC of HIF1α and Sox10 in the forebrain periventricular white matter (Fbr-PWM, marked by dotted lines) of HIFα intact mice **(C)** and HIFα stabilized *(Sox10-Cre:Vhf^m^)* mice **(D)** at P5. **E-F**, IHC of HIF1α and Sox10 in the Fbr-PWM of P10 mice that had been subjected to sham operation **(E)** and hypoxia/ischemia (H/I) injury on P6, i.e. 4 days post-H/I or sham (see Materials and Methods for details). Arrowheads in **A-F** point to HIF1α^+^Sox10^+^ cells. **G**, Western blot of micro-dissected subcortical white matter for HIF1α and canonical HIFα target PKM2 at 6 hours post-H/I or sham. GAPDH, internal loading control. **H**, PKM2 IHC in the Fbr-PWM at P10, 4 days post-HlI or sham. **I1-I3**, IHC and quantification of myelin basic protein (MBP) staining in the Fbr-PWM at P10, 4 days post-HlI or sham. MBP intensity was normalized to that of the contralateral brain hemispheres to the occluded artery of the same mouse. Scale bars, 20 μm.

HIF1α signals became barely detectable in the forebrain white matter by P10 (Fig. 1E), which temporally approximates to the peak of CNS angiogenesis (Harb et al., 2013; Yuen et al., 2014), and remarkably upregulated in mice that had been challenged by hypoxia/ischemia injury at P6 (Fig. 1F) (Chung et al., 2013; Liu et al., 2011; Shen et al., 2010). Our quantification data showed that the density of HIF1α^+^Sox10^+^ cells were significantly increased in P10 H/I-injured forebrain white matter compared with sham controls (4.2 ± 4.2 sham, 312 ± 9.7 H/I, mean ± s.e.m., n=3, t_(4)_ = 29.3 *P* < 0.0001). Consistent with HIF1α histological upregulation, Western blot assays of micro-dissected subcortical white matter demonstrated a substantial elevation of HIF1α and its canonical target gene PKM2 in H/I-injured brain (Fig. 1G) which were also confirmed by PKM2 IHC (Fig. 1H). Concomitant with HIFα activation, myelin staining by myelin basic protein (MBP) showed severe disturbance of developmental myelination in H/I-injured white matter compared with sham control (Fig. 1I1-3). These results demonstrate that HIFα is transiently stabilized in the early postnatal CNS but rapidly downregulated in the second postnatal week and that hypoxia/ischemia injury results in sustained HIFα stabilization in the brain white matter.

### HIFα hyperactivity impairs OPC differentiation but not oligodendrocyte maturation

Pathological hypoxia-ischemia insults caused aberrant HIFα activation in the white matter (Fig. 1F-H) and hypomyelination (Fig. 1I1-3). To determine whether aberrant HIFα stabilization disturbs normal developmental myelination, we induced HIFα hyperactivity by conditionally disrupting the negative regulator VHL (cf Fig. 1B, D). Constitutive *Sox10-Cre:Vhl^fl/fl^* (Sox10:VHL cKO) pups were much smaller than littermate controls (Vhl^fl/+^, or Vhl^fl/fl^) at birth and rarely survived beyond P7. Our prior data show that Sox10-Cre-mediated gene disruption initiates in early OPCs during embryonic CNS development (Zhang et al., 2018a). *Sox10-Cre-* elicited HIFα hyperactivity (cf Fig. 6A) caused severe hypomyelination in the developing postnatal spinal cord (Fig. 2A). The number of CC1^+^ mature OLs (Fig. 2B, C) and the mRNA level of major myelin protein (Fig. 2D) were reduced by ∼40-60% in Sox10:VHL cKO mutants compared with littermate controls. However, the number of PDGFRα^+^ OPCs was unaffected in Sox10:VHL cKO mutants (Fig. 2C). These data suggest that constitutive HIFα activation in OPCs inhibits OPC differentiation and disturbs developmental myelination.

**Figure 2.**
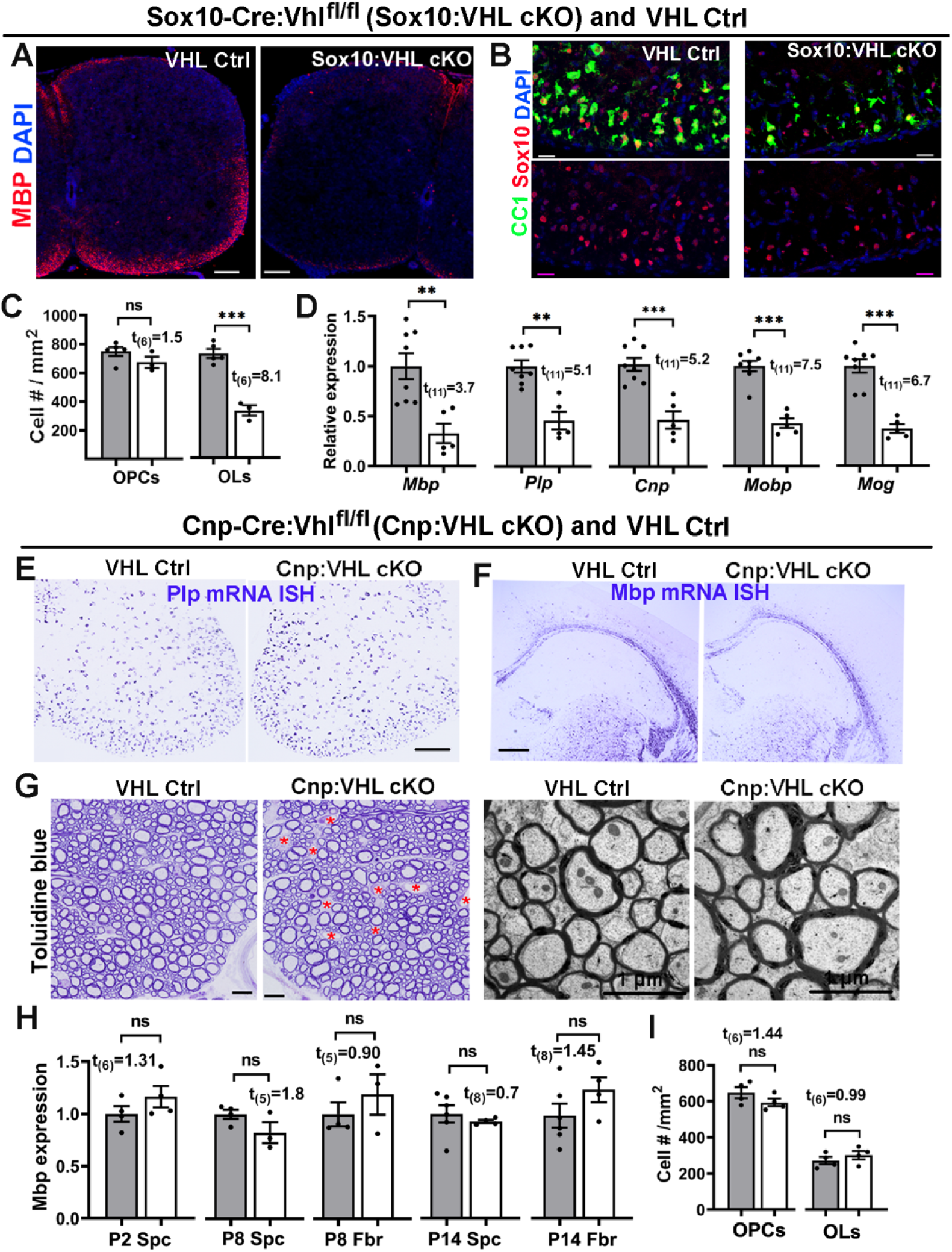
HIFα activation impairs developmental myelination by inhibiting OPC differentiation but not subsequent OL maturation. **A**, Myelin staining of MBP in the spinal cord of *Sox10-Cre:Vhf^m^* (Sox10:VHL cKO) and littermate control *(Vhf’^l^*^+^ or *Vhl*/^fl/fl^) pups at P5. Most Sox10:VHL cKO pups died by the first postnatal week. **B**, representative confocal images of Sox10 (red) and differentiated OL marker CC1 (green) in P5 spinal cord (n=5 Ctrl, 3 cKO). Blue, DAPI nuclear counterstaining. **C**, quantification of Sox10^+^CC1^+^ OLs and Sox10^+^PDGFRa^+^ OPCs (right) in P5 spinal cord (n=5 Ctrl, 3 cKO). **D**, RT-qPCR assay for myelin-specific genes in the spinal cord at P5. *Pip*, proteolipid protein; *Cnp*, 2’,3’-Cyclic-nucleotide 3’-phosphodiesterase; *Mobp*, myelin oligodendrocyte basic protein; *Mog*, myelin oligodendrocyte glycoprotein. **E-F**, mRNA in situ hybridization (ISH) of *Pip* and *Mbp* in the spinal cord **(E)** and forebrain **(F)** of *Cnp-Cre:VhP^m^* (Cnp:VHL cKO) and littermate control *(Cnp-Cre:Vhl^fl+^*, *Vhf’^l^*^+^, or *Vhf’^fl/fl^)* mice at P14. **G**, myelination in the adult spinal cord indicated by toluidine blue staining of semithin sections (left) and transmission electron microscopy (TEM) of ultrathin section (right). Red asterisks in **G** denotes blood vessels, the density of which is elevated in Cnp:VHL cKO mice (Guo et al., 2015). **H**, *Mbp* mRNA levels quantified by RT-qPCR in Cnp:VHL cKO and Ctrl mice at different time points. Spc, spinal cord; Fbr, forebrain. **I**, densities of Sox10^+^CC1^+^ OLs and Sox10^+^PDGFRa^+^ OPCs in P8 spinal cord of Cnp:VHL cKO and Ctrl mice. Scale bars: A, 100μm, B, 20μm, E-F, 200μm, G, 10μm (left), μm (right). Data are presented as mean ± s.e.m.; gray bars are from Ctrl, white bars cKO; two-tailed Student’s *t* test; * p<0.05, ** p<0.01, *** p<0.001, ns, not significant. The above data presentation and statistics are applied to all figures except indicated otherwise).

To determine the effect of HIFα hyperactivity on subsequent oligodendrocyte maturation, we analyzed *Cnp-Cre:Vhl^fl/fl^* (Cnp:VHL cKO) mutants. Previous studies have demonstrated that, compared with *Sox10-Cre, Cnp-Cre* drives gene disruption in later stages of oligodendrocyte development (Moyon et al., 2016; Zhang et al., 2018a; Zhang et al., 2018b). Cnp:VHL cKO mice were viable and indistinguishable from their littermate controls (*Vhl*^fl/fl^, Vh^fl/+^, or *Cnp-Cre:Vhl*^fl/+^) throughout the early postnatal and adult ages. In sharp contrast to Sox10:VHL cKO mice, Cnp:VHL cKO mice had normal levels of myelin gene expression (Fig. 2E, F, H), unperturbed myelination evidenced by toluidine blue myelin staining (Fig. 2G, left) and transmission electron microscopic (TEM) myelination assay (Fig. 2G, right), and normal numbers of OLs and OPCs (Fig. 2I) despite the elevated density of blood vessels in the CNS we recently reported (Guo et al., 2015; Zhang et al., 2020). These data indicate that HIFα hyperactivity in OLs does not affect their maturation. Taken together, our genetic evidences convincingly demonstrate elevated HIFα activity perturbs developmental myelination by inhibiting OPC differentiation but not OL maturation, suggesting a lineage stage-dependent role of HIFα in developmental myelination.

We next used time-conditional and OPC-specific genetic model to circumvent the early death of Sox10:VHL cKO mice and interrogate whether time-conditional HIFα activation in postnatal OPCs impacts their differentiation. Towards this end, we generated *Pdgfrα-CreER^T2^:Vhl^fl/fl^* (PDGFRa:VHL cKO) mutants. Our data showed that OPC-specific HIFα activation during neonatal ages (Fig. 3A-B) significantly reduced the expression of genes encoding major myelin proteins (Fig. 3C) and decreased the densities of mature OLs but not OPCs in the spinal cord (Fig. 3D). IHC of myelin protein PLP demonstrated a marked hypomyelination (Fig. 3E). Further analysis of myelination by transmission electron microscopy (Fig. 3F) demonstrated that the density of myelinated axons was reduced by ∼42% in the ventral spinal white matter of PDGFRa:VHL cKO mice compared with littermate control (Fig. 3G), which is consistent with histological myelin protein assays (Fig. 3F). To assess the long­term effect of HIFα activation, we induced HIFα activation by tamoxifen injections to neonatal mice (Fig. 3H-I) and analyzed oligodendroglial phenotypes in the adult CNS (P60-P70). We found that the number of OLs but not OPCs was persistently reduced in the corpus callosum of PDGFRa:VHL cKO compared with control mice (Fig. 3J), suggesting that neonatal HIFα activation inhibits OPC differentiation in the adult CNS. Given that virtually all axons in the optic nerve are myelinated in the adult rodents (Bercury and Macklin, 2015), we then analyzed myelinated axons visualized by toluidine blue myelin staining of semithin resin sections (Fig. 3K1-K4). Our quantification demonstrated a ∼44% reduction in the density of myelinated axons in the optic nerve of adult PDGFRa:VHL cKO mutants compared with age-matched fully myelinated control optic nerves (Fig. 3K5). Together, our data suggest that HIFα activation inhibits long-term OPC differentiation and results in hypomyelination throughout postnatal CNS development.

**Figure 3.**
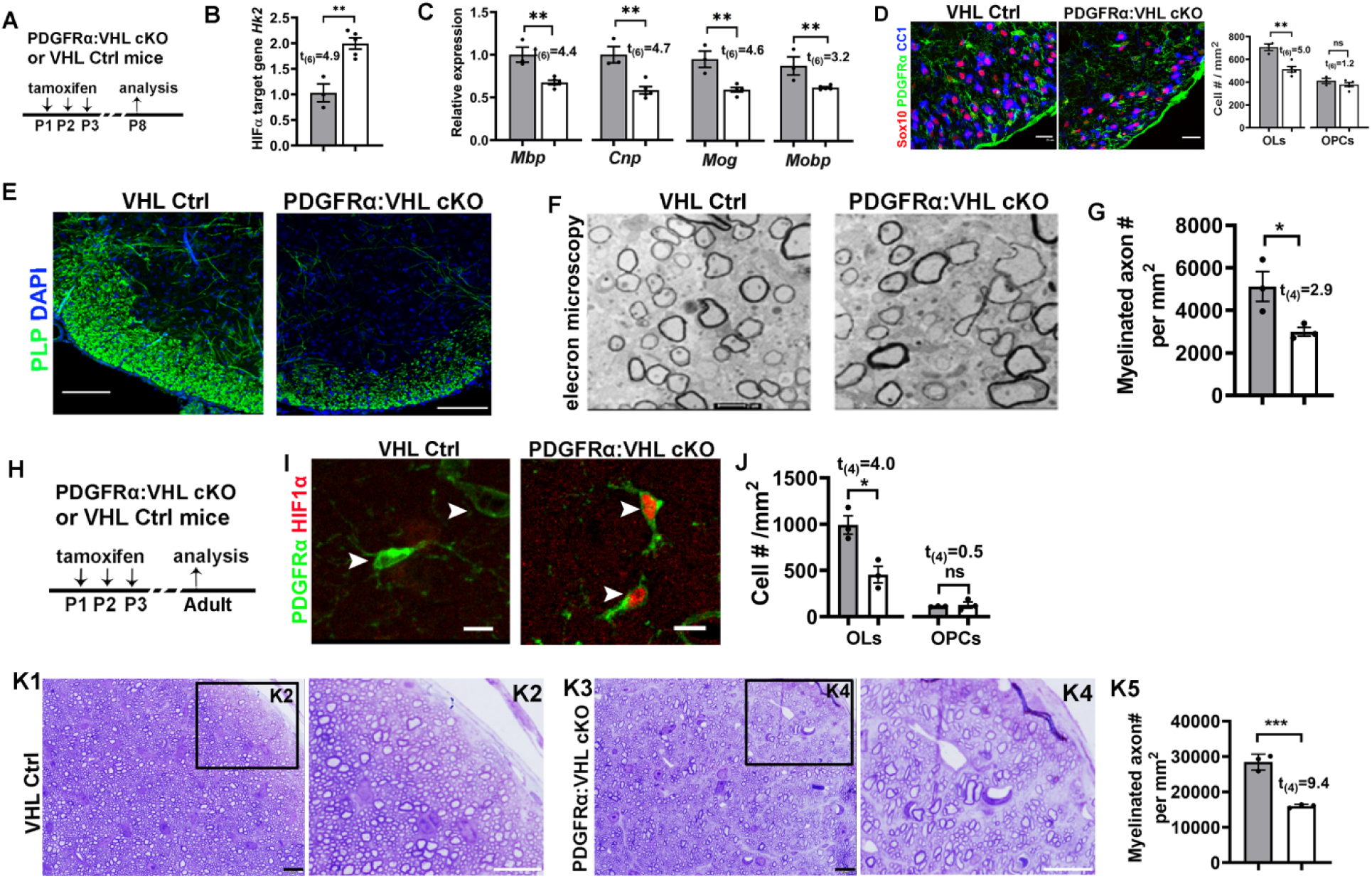
Inducible HIFα stabilization in postnatal OPCs inhibits OPC differentiation. **A**, experimental design for B-**F**. PDGFRccVHL cKO *(Pdgfrα-CreER^T2^:Vhf^m^)* and littermate control (Vh/^l/fl^) mice were treated with tamoxifen intraperitoneally at P1, P2, and P3, and analyzed at P8. **B**, RT-qPCR quantification of HIFα target gene hexokinase 2 *Hk2* in the spinal cord. **C**, expression of major myelin proteins in the spinal cord quantified by RT-qPCR. **D**, representative confocal images and the density of Sox10^+^/CC1^+^ OLs and Sox10^+^/PDGFRα^+^ OPCs. **E**, IHC of PLP showing myelination in the spinal cord ventral white matter. **F-G**, representative TEM images **(F)** and the density of myelinated axons **(G)** in the spinal cord ventral white mater. **H**, experimental design for **I-K**. PDGFRα:VHL cKO and littermate control mice were treated with tamoxifen intraperitoneally at P1, P2, and P3, and analyzed at P60-70. **I**, representative confocal images showing HIF1α stabilization in PDGFRα^+^ OPCs (arrowheads) of PDGFRα:VHL cKO mice. **J**, densities of CC1^+^ mature OLs and PDGFRα^+^ OPCs in the corpus callosum. K1-K4, toluidine blue staining for myelin on semithin (500nm) resin sections showing hypomyelination in the optic nerve of adult PDGFRα:VHL cKO mice. Boxed areas in **K1** and **K3** were shown at higher magnification images in **K2** and **K4**, respectively. **K5**, density of myelinated axons in the adult optic nerves. Scale bars, D, 25μm; E, 100μm; F, 0.5 μm; I-K, 10 μm.

Taken together, our *in vivo* data derived from three different genetic mouse models suggest that HIFα hyperactivity disturbs developmental myelination by inhibiting OPC differentiation.

### Disrupting HIFα in OPCs but not OLs delays developmental myelination

Transient HIFα stabilization (Fig. 1A-E) suggests that HIFα may regulate OPC differentiation and/or myelination. To define the role of physiological HIFα stabilization in developmental myelination, we analyzed oligodendroglial phenotypes in HIFα cKO genetic mice. Our extensive analysis of HIF1α or HIF2α conditionally knockout mice driven by *Pdgfrα-CreER^T2^*and *Cnp-Cre* revealed no phenotypic abnormalities of the oligodendroglial lineage, suggesting functional redundancy between HIF1α and HIF2α. We therefore generated HIF1α and HIF2α double conditional knockout (referred to as HIFα cKO).

To determine the role of HIFα in OPC differentiation, we used *Pdgfrα-CreER*^T2^:*HiF1α*^fl/fl^:*Hif2α*^fl/fl^ (PDGFRα:HIFα cKO) transgenic mice. Tamoxifen was administered to PDGFRα:HIFα cKO and littermate control (*Hif1α*^fl/fl^:*Hif2α*^fl/fl^) pups at P1, P2, and P3 and the CNS tissue was analyzed at P8. Our previous data show that this tamoxifen paradigm induces >85% of gene cKO efficiency in OPCs (Zhang et al., 2018a; Zhang et al., 2018b). The density of differentiated OLs (CC1^+^Olig2^+^) but not OPCs (PDGFRα^+^Olig2^+^) was significantly decreased in the spinal cord white matter of PDGFRα:HIFα cKO mice compared with HIFα controls (Fig. 4A-B), suggesting that HIFα regulates OPC differentiation and is dispensable for OPC population expansion. In agreement of impaired OPC differentiation, the mRNA levels of major myelin protein *(Mbp* and *Plp)* and mature OL-enriched protein (Opalin and QK, aka CC1) were significantly reduced in PDGFRα:HIFα cKO mice compared with littermate controls (Fig. 4C). In the forebrain, developmental myelination in the periventricular white matter was delayed in PDGFRα:HIFα cKO mice at P8 (Fig. 4D) and became indistinguishable from that in littermate control mice at P14 (Fig. 4E). These data suggest that HIFα regulates the timing of developmental myelination.

**Figure 4.**
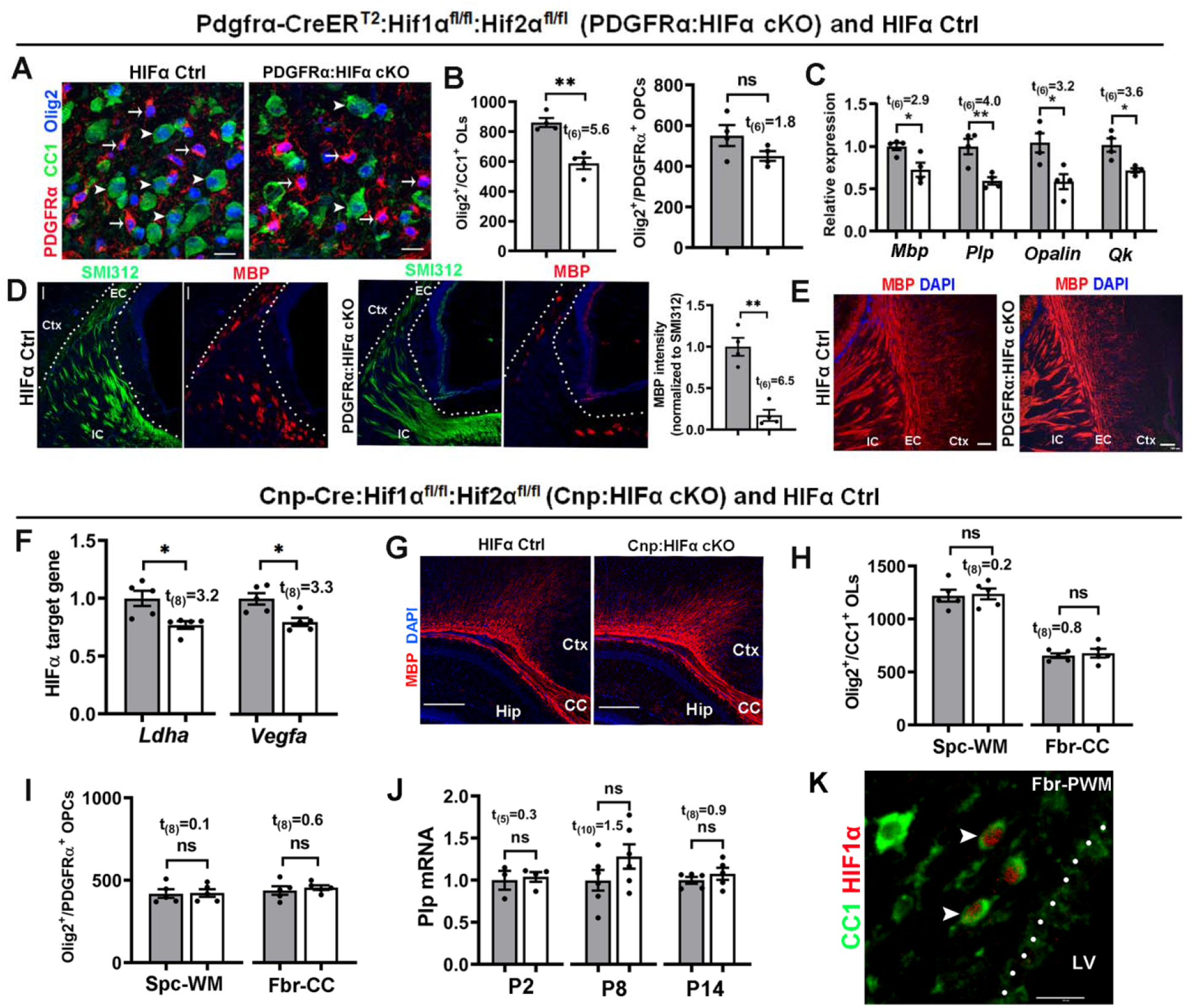
HIFα inactivation transiently delays developmental myelination by controlling OPC differentiation but not subsequent OL maturation. **A-E**, PDGFRα:HIFα cKO and littermate control (HIF1α^fllfl^:HIF2α^fllfl^) mice were treated with tamoxifen at P1, P2, and P3, and analyzed at P8 (n=4 each group) and P14 (n=3 each group). **A-B**, IHC **(A)** and quantification **(B)** of Olig2^+^CC1^+^ OLs and Olig2^+^PDGFRα^+^ OPCs in the spinal cord. **C**, relative expression of myelin protein genes of *Mbp* and *Plp* and mature OL-specific gene of *Opalin* and quake *(Qk*, aka *CC1)* in the forebrain measured by real-time quantitative PCR (RT-qPCR). **D**, IHC of myelin marker MBP and pan-axonal marker SMI312 in forebrain periventricular white matter (left, marked by dotted area) and quantification (right) at P8. EC, external capsule, IC, internal capsule, Ctx, cortex. **E**, IHC of MBP in forebrain periventricular white matter at P14 (tamoxifen at P1, P2, and P3). **F-J**, Cnp:HIFα cKO and littermate control (HIF1α^fllfl^:HIF2α^fllfl^) mice were analyzed at P2, P8, and P14. **F**, RT-qPCR quantification of HIFα target genes *Ldha* and *Vegfa* in P8 spinal cord. **G**, myelin staining by MBP in the forebrain. CC, corpus callosum, Ctx, cortex, Hip, hippocampus. **H-I**, densities (#/mm^2^) of Olig2^+^CC1^+^ OLs **(H)** and Olig2^+^PDGFRα^+^ OPCs **(I)** at P14. Spc-WM: spinal cord white matter, Fbr-CC: forebrain corpus callosum. **J**, RT-qPCR assay of exon3b-containing Plp mRNA, which is specific to mature OLs, in the spinal cord at different time points. **K**, IHC of CC1 and HIF1α in P8 forebrain periventricular white matter (Fbr-PWM). Arrowheads point to double positive cells. LV, lateral ventricle. Scale bar, A, K, 10um; D, 20um; E, 100um; G, 250um.

To determine the role of HIFα in oligodendroglial maturation, we used *Cnp-Cre:Hif1α*^fl/fl^:*Hif2α*^fl/fl^ (Cnp:HIFα cKO) and littermate control (*Hif1α*^fl/fl^:*Hif2α*^fl/fl^) mice. Cnp:HIFα cKO mice were born at expected Mendelian ratios and displayed no behavioral abnormalities during postnatal development and throughout adult ages. The expression of canonical HIFα target gene *Ldha* and *Vegfa* (Sharp and Bernaudin, 2004) was significantly reduced in Cnp:HIFα cKO mice (Fig. 4F), thus validating the disruption of HIFα function. Unexpectedly, we found no significant difference in developmental myelination (Fig. 4G), the number of CC1^+^OLig2^+^ mature OLsand myelin gene expression (Fig. 4H) and PDGFRα^+^Olig2^+^ OPCs (Fig. 4I), and myelin gene expression (Fig. 4J) between Cnp:HIFα cKO and littermate controls mice at different ages (P2, P8, and P14). Double IHC showed that some CC1^+^ OLs were positive for HIF1α in the periventricular white matter of the brain at P8 (Fig. 4K, arrowheads) but not at P14 (not shown). These data indicate that HIFα plays a minor role in oligodendrocyte maturation. Collectively, the contrast phenotypes derived from PDGFRα:HIFα cKO vs Cnp:HIFα cKO mice support a working model that HIFα controls the timing of developmental myelination by regulating the differentiation of OPCs into OLs but not subsequent oligodendrocyte maturation.

### Disrupting oligodendroglial HIFα does not affect axonal integrity or cell survival

HIFα is required for neuronal survival, as HIFα deficiency in Nestin-expressing neural cells results in massive neuronal death and neurodegeneration (Tomita et al., 2003). Recent data reported that HIFα deficiency in Olig1-expressing neural cells leads to neuronal death and widespread axonal injury in the corpus callosum at the perinatal ages (Yuen et al., 2014). To determine whether OPC or OL-specific HIFα deficiency causes axonal damage and cell death, we employed IHC of SMI32 monoclonal antibody and cleaved Caspase 3 (CC3) antibody.

SMI32 labels functionally compromised axons (Soulika et al., 2009; Yuen et al., 2014) and a subset of pyramidal neuronal cell bodies and dendrites during normal development (Campbell and Morrison, 1989; Haynes et al., 2005; Voelker et al., 2004) (Fig. 5A, Ctx). We did not observe SMI32 immunoreactive signals in the corpus callosum of Cnp:HIFα cKO (Fig. 5A) and PDGFRα:HIFα cKO (Fig. 5C) mice compared with respective littermate controls. In contrast, many tightly packed axons running through the corpus callosum were clearly labeled by the pan-axonal marker SMI312 (Fig. 5B, D). Furthermore, the densities of CC3^+^ apoptotic cells and CC3^+^/Sox10^+^ apoptotic oligodendroglial lineage cells were statistically indistinguishable between PDGFRα (or Cnp):HIFα cKO mutants and their respective littermate controls (Fig. 5E-G). We employed TEM to assess the structural integrity of axons in the corpus callosum of PDGFRα:HIFα cKO mice at P8, a time point that few axons are myelinated yet. The density of axons with diameter ≥ 400nm, which will be myelinated by oligodendrocyte during postnatal development (Hines et al., 2015; Lee et al., 2012), was similar between PDGFRα:HIFα cKO (458,401 ± 59,957 axons/mm^2^) and littermate control (485,558 ± 38,780 axons/mm^2^, *t*_(4)_ = 0.4, p = 0.72), suggesting that axons in HIFα cKO mice are not subjected to degeneration. We did not find visible morphological differences in axonal structure (Fig. 5H-I, Ax) and axonal mitochondria (Fig. 5H-I, Mt) between PDGFRα:HIFα cKO and littermate control mice. Thus, disrupting oligodendroglial HIFα does not affect neuronal cell survival and axonal integrity in the white matter in the early postnatal CNS.

**Figure 5.**
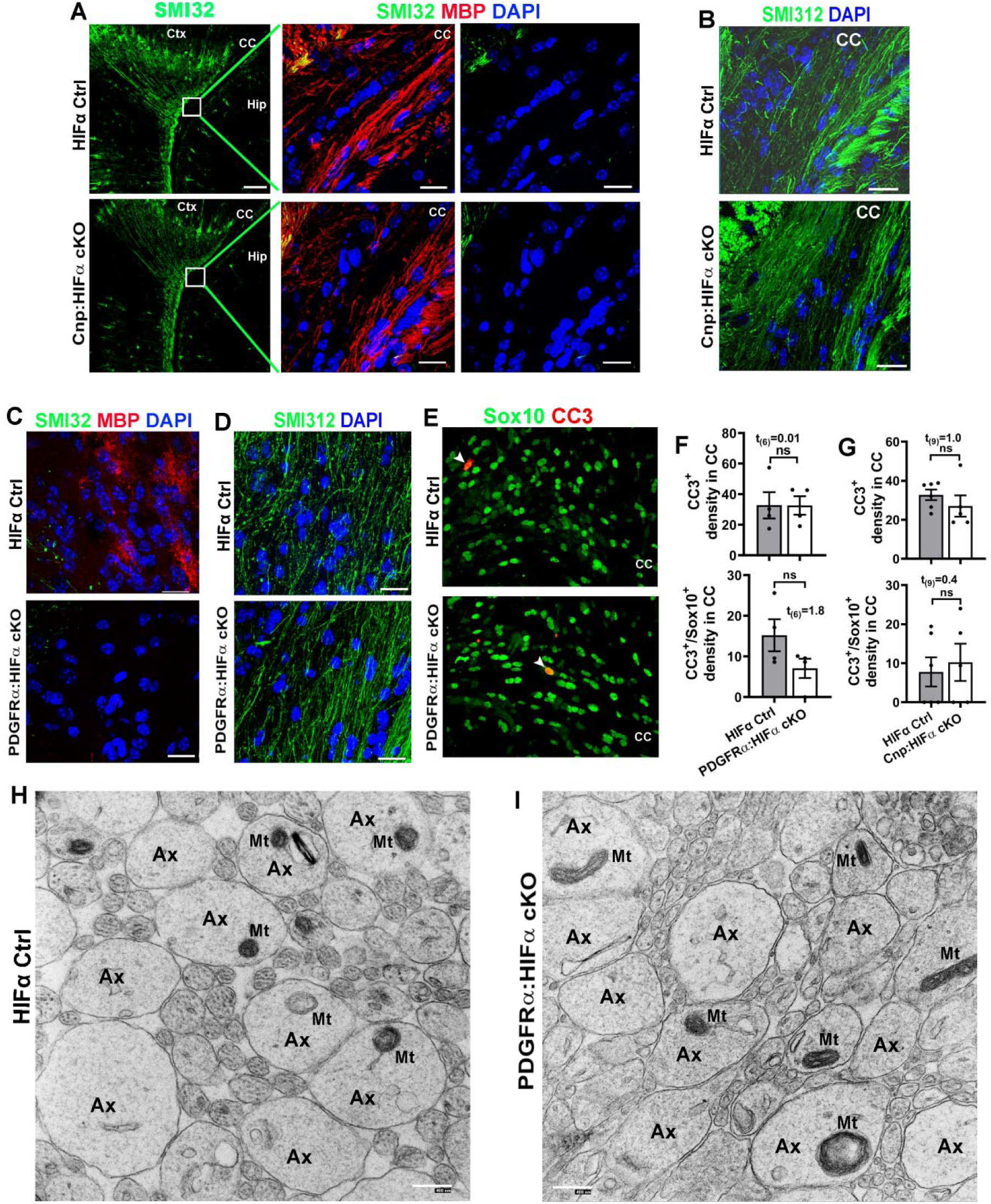
Oligodendroglial HIFα is dispensable for axonal integrity or cell death. **A**, immunostaining images of P14 forebrain using SMI32, a monoclonal antibody recognizing the non-phosphorylated neurofilament proteins of heavy chain (NFH), which has been reported labeling injured axons and a subset of pyramidal neuron bodies and dendrites. Boxed areas show the corpus callosum (CC) at higher magnification images of SMI32 and MBP in the right panels. Ctx, cortex, Hip, hippocampus. Note the absence of SMI32-positive signals in the CC of both Cnp:HIFα cKO and Ctrl mice. **B**, immunostaining images of P14 CC using SMI312, a monoclonal antibody recognizing both phosphorylated and non-phosphorylated NFH (a pan-axonal marker). **C-D**, IHC of SMI32/MBP **(C)** and SMI312 **(D)** in the CC of P8 non-Cre Ctrl and PDGFRcrHIFα cKO and littermate control (HIF1α^fl/fl^:HIF2α^fl/fl^) mice (tamoxifen injection at P1, P2, and P3). **E-F**, representative confocal images and quantification of cells positive for cleaved caspase 3 (CC3) and/or pan-oligodendroglial lineage marker Sox10 in P8 CC of PDGFRcrHIFα cKO and control mice. Arrowheads point to CC3^+^/Sox10^+^ cells. **G**, quantification of CC3^+^ and CC3^+^/Sox10^+^ cells in P14 CC of Cnp:HIFα cKO and control mice. **H-I**, representative TEM images showing the cross-sections of axons in the corpus callosum of HIFα Ctrl **(I)** and cKO **(H)** mice at P8 (tamoxifen injections at P1, P2, and P3) when myelination is barely detectable at the TEM level. Note that the morphology of axons (Ax) and axonal mitochondria (Mt) were indistinguishable between HIFα Ctrl and cKO mice. Scale bar, A, 100μm (low magnification), 20μm (high magnification); B-E, 20μm, H-I, 400nm.

### Oligodendroglial HIFα plays a minor role, if any, in regulating Wnt/p-catenin activity

Previous study reported that aberrant HIFα stabilization in OPCs activates Wnt7alWnt7b expression and canonical Wnt signaling (i.e. Wnt/β-catenin) (Yuen et al., 2014). The unbiased RNA-seq analysis of *Sox10-Cre:Vhl*^fl/fl^ mice prompted us to revisit HIFα-Wnt regulation in oligodendroglial lineage cells (Fig. 6A-E). Sox10-Cre-mediated VHL cKO markedly elevated HIFα activity, as evidenced by increased expression of canonical HIFα target genes *HK2, Ldha, Glutl, Bnip3*, and *Vegfa* (Fig. 6A). In stark contrast to HIFα activation, there were no significant changes in the expression of Wnt genes *Wnt7a* and *Wnt7b* and canonical Wnt signaling target genes *Axin2* and *Naked1* (Fig. 6B). The KEGG pathway of HIF-1 signaling and the GO term of response to hypoxia were significantly overrepresented amongst 255 upregulated genes (Fig. 6C, D) whereas the GO terms of oligodendrocyte differentiation and sterol biosynthesis were significantly overrepresented amongst 351 downregulated genes (Fig. 6E), which is in agreement with elevated HIFα signaling and diminished OPC differentiation in the CNS of Sox10:VHL cKO mice we documented in Fig. 2A-D. However, we did not find overrepresentation of Wnt-related KEGG pathways or GO terms amongst the differentially regulated genes. The unbiased analysis of gene regulation at the transcriptomic level strongly suggests that oligodendroglial HIFα stabilization plays a minor role in regulating canonical Wnt signaling.

**Figure 6.**
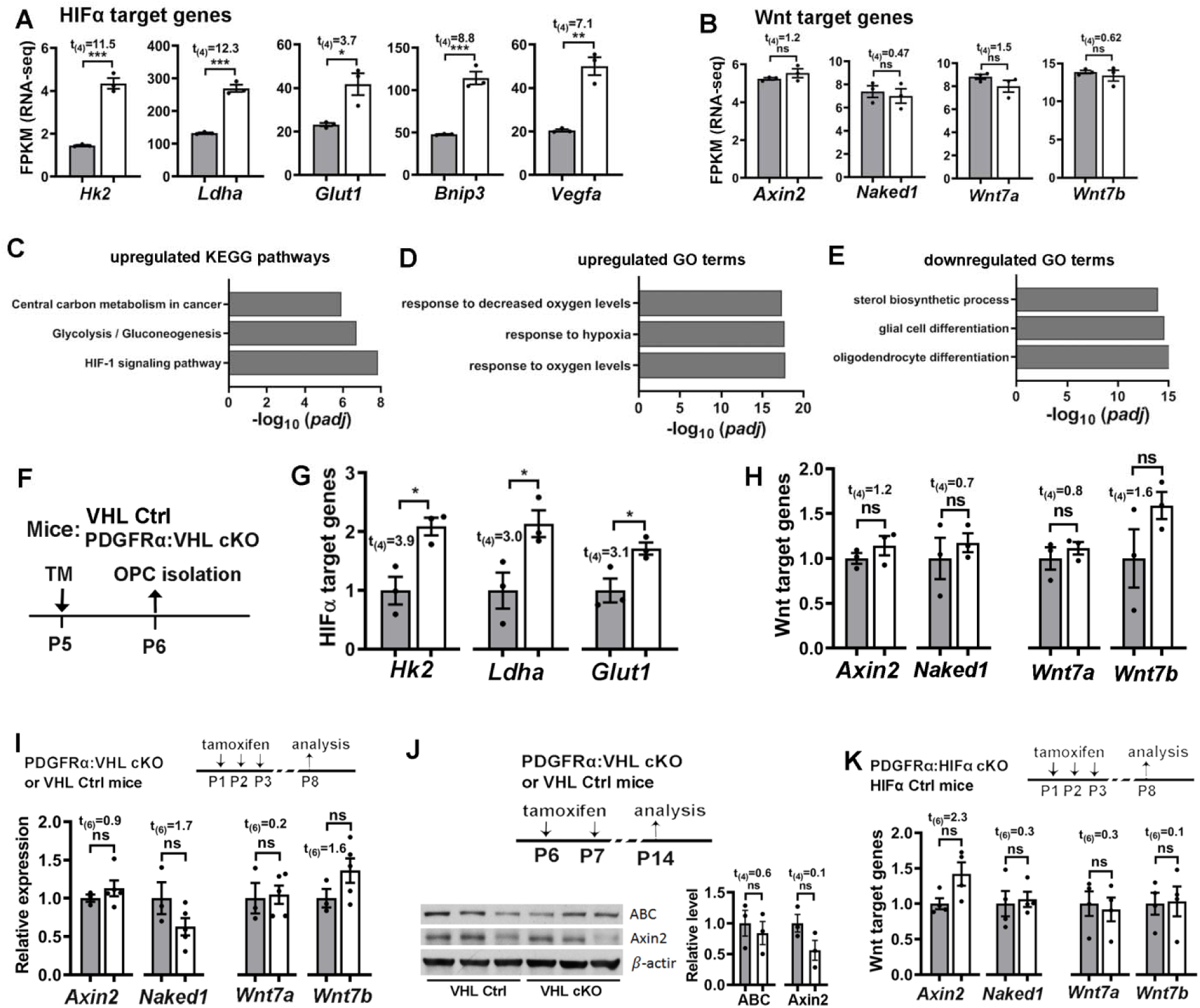
Stabilizing HIFα activates HIFα-mediated signaling but does not perturb Wnt/β-catenin signaling in the oligodendroglial lineage and in the CNS. **A-B**, relative expression of HIFα and Wnt target genes from unbiased RNA-seq analysis. FPKM, fragments per kilobase of transcript per million mapped reads, a normalized measurement of gene expression. The RNA was prepared from the spinal cord of P5 *Sox10-Cre:Vhl*^fl/fl^ mice (n=3, white bars) and littermate control (*Vhl*^fl/+^ or *Vhl^fl/fl^*) (n=3, gray bars) mice. **C-E**, Top 3 significant enriched KEGG pathway and gene ontology (GO) terms among 255 upregulated and 351 downregulated genes derived from unbiased RNA-seq. Note that HIFα signaling pathway **(C)** and response to hypoxia **(D)** were significantly upregulated, validating HIFα hyperactivation in *Sox10-Cre:Vhl^m^* mice and that oligodendroglial cell differentiation was significantly downregulated **(E)**, which is agreement with the data of Fig. 2A-D. **F**, experimental design for G and H. PDGFRα:VHL cKO (n=3, white bars) and littermate control (Vh/^l/fl^, n=3, gray bars) mice were treated with tamoxifen (TM) at P5, and forebrain OPCs at P6 were acutely purified by MACS. **G-H**, RT-qPCR quantification of HIFα target gene **(G)** and Wnt target gene **(H)** expression in purified OPCs. **I**, expression of Wnt target gene *(Axin2* and *Naked1)* and *Wnt7a/7b* in P8 spinal cord (n=3 Ctrl, gray bars, 5 PDGFRα cKO, white bars) quantified by RT-qPCR. **J**, Western blot images and relative expression (normalized to the internal loading control β-actin) of the active form of β-catenin (ABC) and Axin2 in P14 forebrain (n=3 each group). **K**, expression of Wnt target genes *(Axin2* and *Naked1)* and *Wnt7a/7b* in P8 spinal cord measured by RT-qPCR (n=4 each group).

To strengthen our conclusion derived from RNA-seq, we analyzed OPCs isolated from PDGFRα:VHL cKO mice. Tamoxifen was injected to PDGFRα:VHL cKO and littermate control mice at P5 and brain OPCs were acutely purified at P6 (Fig. 6F) by magnetic activated cell sorting (MACS). As shown in Fig. 6G, HIFα was indeed functionally stabilized in purified VHL-disrupted OPCs, as evidenced by the elevated expression of HIFα target gene *Hk2, Ldha*, and *Glut1.* However, there were no significant changes in the expression of canonical Wnt/β-catenin signaling target genes *Axin2* and *Naked1* and Wnt ligands *Wnt7a* and *Wnt7b* in VHL-disrupted OPCs compared with VHL-intact OPCs (Fig. 6H), suggesting that genetically stabilizing HIFα does not perturb the activity of autocrine Wntl3-catenin or the expression of Wnt7a and Wnt7b in OPCs.

To corroborate our findings, stabilizing HIFα in PDGFRα^+^ OPCs did not affect the activity of Wnt/β-catenin signaling in the spinal cord at P8 (tamoxifen at P1, P2, and P3) and in the forebrain at P14 (tamoxifen at P6 and P7) of PDGFRα:VHL cKO mice compared with littermate controls, which was supported by the unaltered expression of canonical Wnt target genes at the mRNA and protein levels (Fig. 6I-J). We also found that the expression of Axin2, Naked1, Wnt7a, and Wnt7b was not perturbed in PDGFRα:HIFα cKO mice compared with littermate controls (Fig. 6K). Thus, HIFα does not regulate canonical Wnt signaling activity or Wnt7a/Wnt7b expression in the oligodendroglial lineage and in the CNS.

### Blocking oligodendroglial-derived Wnt signaling does not affect HIFα hyperactivation-elicited inhibition of OPC differentiation and myelination

That non-perturbation of Wnt signaling in HIFα-stabilized OPCs led us to hypothesize that HIFα hyperactivation inhibits myelination in a manner independent of autocrine Wnt activation in OPCs. To test our hypothesis, we stabilized HIFα function and simultaneously blocked oligodendroglial-derived autocrine Wnt signaling.

Previous studies demonstrated that disrupting WLS blocks the secretion of Wnt ligands from Wnt-producing cells and inhibits intracellular Wnt signaling activity in Wnt-receiving cells (Banziger et al., 2006; Bartscherer et al., 2006). To determine whether WLS deficiency blocks autocrine Wnt/β-catenin activity in OPCs, we knocked down WLS in Wnt3a-expressing primary OPCs (Fig. 7A-C). We chose Wnt3a because it is a typical Wnt ligand that activates the canonical Wnt signaling in Wnt-receiving cells. Wnt3a expression in OPCs significantly increased Wnt3a concentration and simultaneous WLS knockdown abrogated Wnt3a elevation in the culture medium of primary OPCs (Fig. 7D), suggesting that WLS is required for Wnt3a secretion from OPCs. We found that the activity of the canonical Wnt signaling, as assessed by Wnt target genes *Axin2* and *Sp5*, was upregulated in Wnt3a-expressing OPCs but blocked in Wnt3a-expressing/WLS-deficient OPCs (Fig. 7E, F). These results demonstrate that disrupting WLS blocks OPC-derived autocrine Wnt/β-catenin signaling activity.

**Figure 7.**
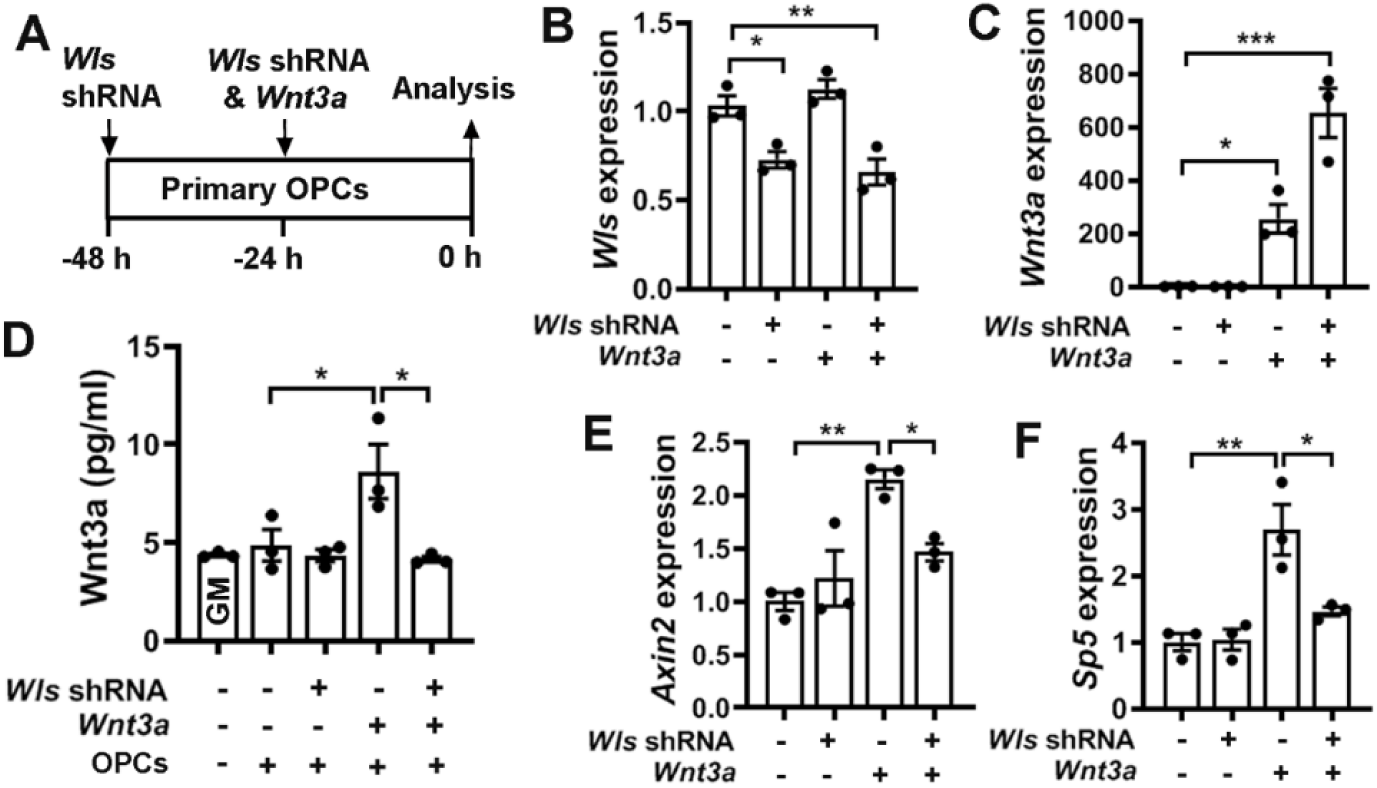
WLS deficiency inhibits Wnt3a secretion and blocks autocrine Wnt/p-catenin signaling activity in OPCs. **A**, experimental design of cell transfection of Wls shRNA and Wnt3a-expressing plasmids to primary brain OPC maintained in the growth medium. **B-C**, RT-qPCR assay of *Wls* **(B)** and *Wnt3a* **(C)** expression in transfected primary OPCs. One-way ANOVA followed by Tukey’s multiple comparisons, *F*_(3, 8)_ = 15.39, *P* = 0.001 Wls, *F*_(3, 8)_ = 32.81, *P* < 0.0001 *Wnt3a.* **D**, ELISA quantification of Wnt3a in the growth medium in the absence or presence of OPCs transfected by Wls shRNA and/or Wnt3a. One-way ANOVA followed by Tukey’s multiple comparisons, *** *P* < 0.001, ns not significant. *F*_(4, 10)_ = 6.70, *P* < 0.0069. Note that Wnt3a concentration in the GM in the presence of primary OPCs is statistically indistinguishable from that in the GM alone, suggesting that intact primary OPCs do not secrete Wnt3a. **E-F**, RT-qPCR quantification of Wnt target genes *Axin2* and *Sp5* in OPCs transfected with Wls shRNA and/or Wnt3a. One-way ANOVA followed by Tukey’s multiple comparisons, *F*_(3, 8)_ = 11.06, *P* = 0.0032 *Axin2*, *F*_(3,8)_ = 13.25, *P* = 0.0018 *Sp5.*

Genetically disrupting WLS in PDGFRα^+^ OPCs had no rescuing effects on HIFα hyperactivation-elicited hypomyelination (Fig. 8A-C), impaired OPC differentiation (Fig. 8D-E), and diminished myelin gene expression (Fig. 8F) in PDGFRα:VHL/WLS cKO mice, suggesting that autocrine Wnt signaling is unlikely a downstream pathway in mediating HIFα stabilization-induced inhibition of OPC differentiation. To corroborate this conclusion, we used a different inducible genetic model, *Sox10-CreER*^T2^:*Vhl*^fl/fl^:*Wls*^fl/fl^ (iSox10:VHL/WLS cKO) (Fig. 9A). Conditionally stabilizing HIFα in Sox10-expressing oligodendroglial lineage cells resulted in impaired OPC differentiation and hypomyelination in iSox10:VHL cKO mice, however, simultaneous WLS cKO did not affect the degree of inhibited OPC differentiation and hypomyelination in iSox10:VHL/WLS cKO mice (Fig. 9B-E). We further demonstrated that disrupting WLS alone in Sox10-expressing oligodendroglial lineage cells had no detectable effect on normal OPC differentiation and developmental myelination in *Sox10-CreER*^T2^:*Wls*^fl/fl^(iSox10:WLS cKO) mice compared with non-Cre littermate control mice (Fig. 10). Collectively, our results derived from cell-specific genetic approaches suggest that autocrine Wnt/α-catenin signaling plays a dispensable role in HIFα-regulated OPC differentiation and myelination.

**Figure 8.**
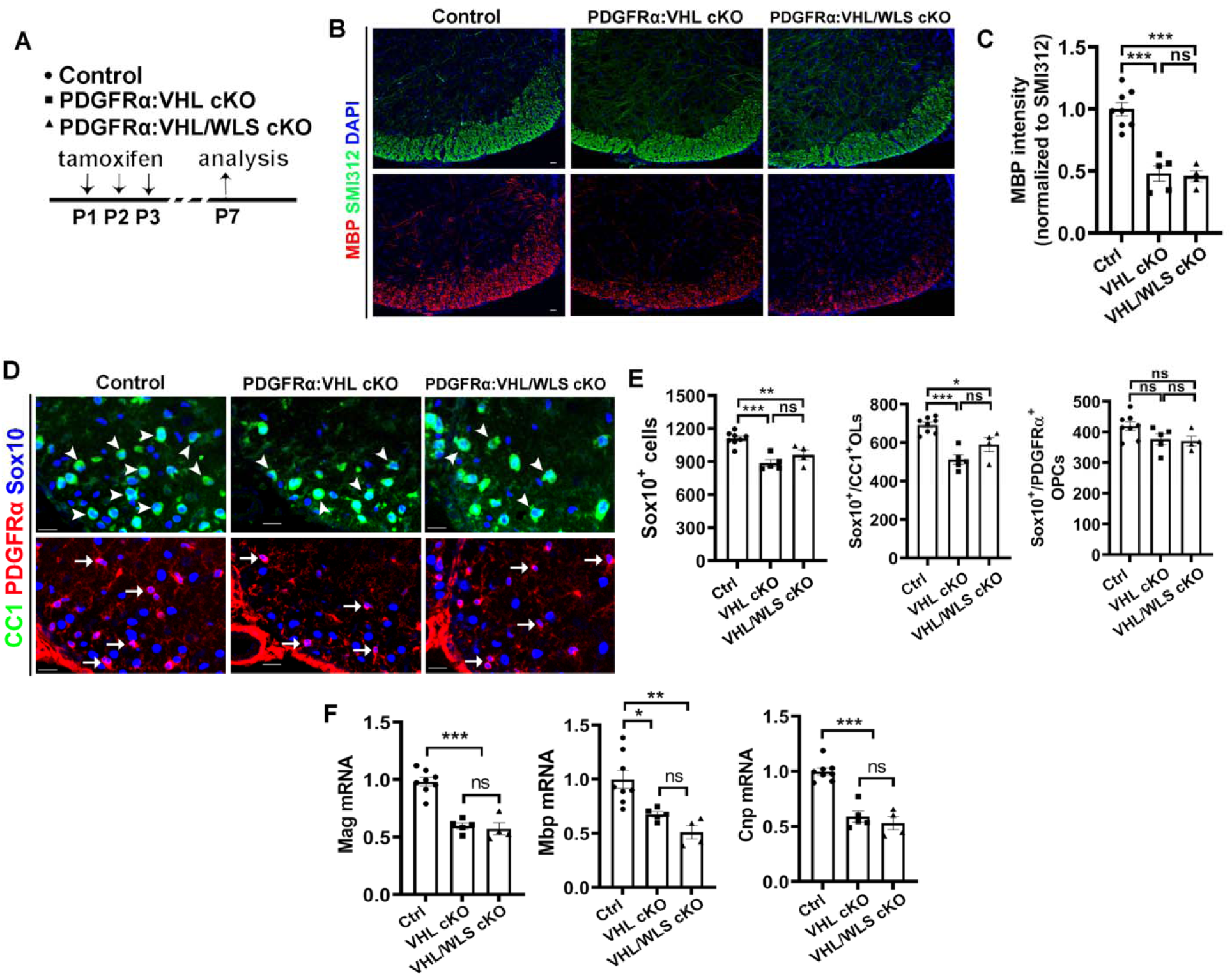
Disrupting WLS in PDGFRα-expressing OPCs does not affect the degree of inhibited OPC differentiation and hypomyelination elicited by HIFα hyperactivation. **A**, experimental design for **B-F**. PDGFRα:VHL cKO, *PDGFRα-CreER^2^:Vhf^m^* (n=5); PDGFRα:VHL/WLS cKO, *Pdgfrα-CreER^12^:* *Vhl*/^fl/fl^:Wls^fl/fl^ (n=4); non-Cre Ctrl carrying *VhH* and/or Wls^fl/fl^ (n=8). **B-C**, representative confocal images **(B)** and quantification **(C)** of myelination by MBP staining. The signal of SMI312, pan-axonal marker is indistinguishable among each group and used as an internal control of MBP quantification. One-way ANOVA followed by Tukey’s multiple comparisons, *F*_(2, 14)_=31.93 P< 0.0001. **D-E**, representative confocal images **(D)** and quantification **(E)** of Sox10^+^/CC1^+^ differentiated OLs (**D**, arrowheads), Sox10^+^/PDGFRα^+^ OPCs (**D**, arrows), and Sox10^+^ oligodendroglial lineage cells in the spinal cord. One-way ANOVA followed by Tukey’s multiple comparisons, *F*_(2, 14)_ = 17.34 *P =* 0.0002 Sox10^+^, *F*_(2, 14)_ = 18.44 *P =* 0.0001 Sox10^+^/CC1^+^, F(_2, 14_) = 2.967 *P =* 0.0843 Sox10^+^/PDGFRα^+^. F, RT-qPCR quantification of mature OL-enriched genes in the spinal cord. One-way ANOVA followed by Tukey’s multiple comparisons, *F*_(2, 14)_ = 11.53 *P =* 0.0011 *Mbp*, *F*_(2, 14)_ = 39.88 *P <* 0.0001 *Mag*, *F*_(2, 14)_ = 38.78 *P <* 0.0001 *Cnp.* Scale bars=20μm in **B** and **D**.

**Figure 9.**
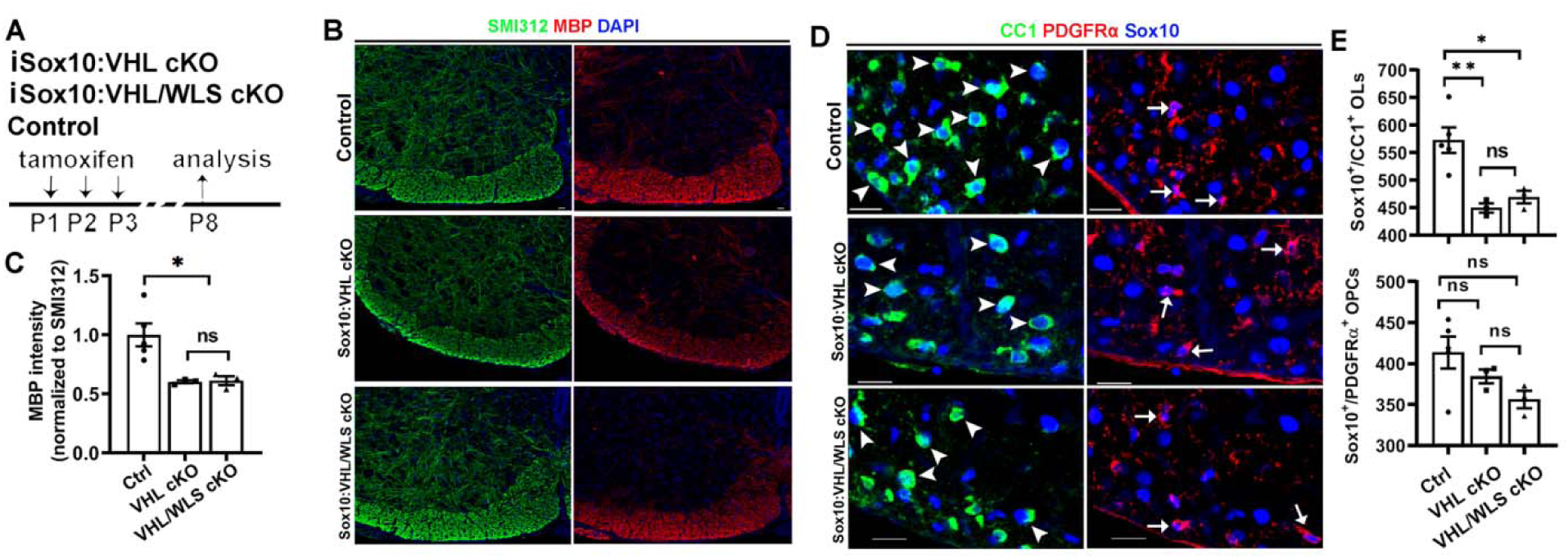
Disrupting WLS in Sox10-expressing oligodendroglial lineage cells does not affect HIFα hyperactivation-elicited inhibition of OPC differentiation and hypomyelination. **A**, experimental design for **B-E**. tamoxifen inducible *Sox10-CreER^T2^:Vhl*^fl/fl^ (iSox10:VHL cKO n=3); *Sox10-CreER^72^: Vhf^m^:Wls^m^* (iSox10:VHLlWLS cKO, n=3); non-Cre Ctrl carrying *Vhl*/^fl/fl^ andlor Wls^fllfl^ (n=5). **B-C**, representative confocal images **(B)** and quantification **(C)** of myelination by MBP staining of the spinal cord. SMI312 signal was used as an internal control of MBP quantification. One-way ANOVA followed by Tukey’s multiple comparisons, *F*_(2, 8)_= 8.484 *P =* 0.0105. **D-E**, representative confocal images **(D)** and quantification **(E)** of Sox10^+^lCC1^+^ differentiated OLs (**D**, arrowheads) and Sox10^+^lPDGFRα^+^ OPCs (**D**, arrows) in the spinal cord. One-way ANOVA followed by Tukey’s multiple comparisons, *F*_(2, 8)_ = 11.90 *P =* 0.004 Sox10^+^lCC1^+^, F(_2, 14_) = 2.880 *P =* 0.1143 Sox10^+^lPDGFRα^+^. Scale bars=20μm in **B** and **D**.

**Figure 10.**
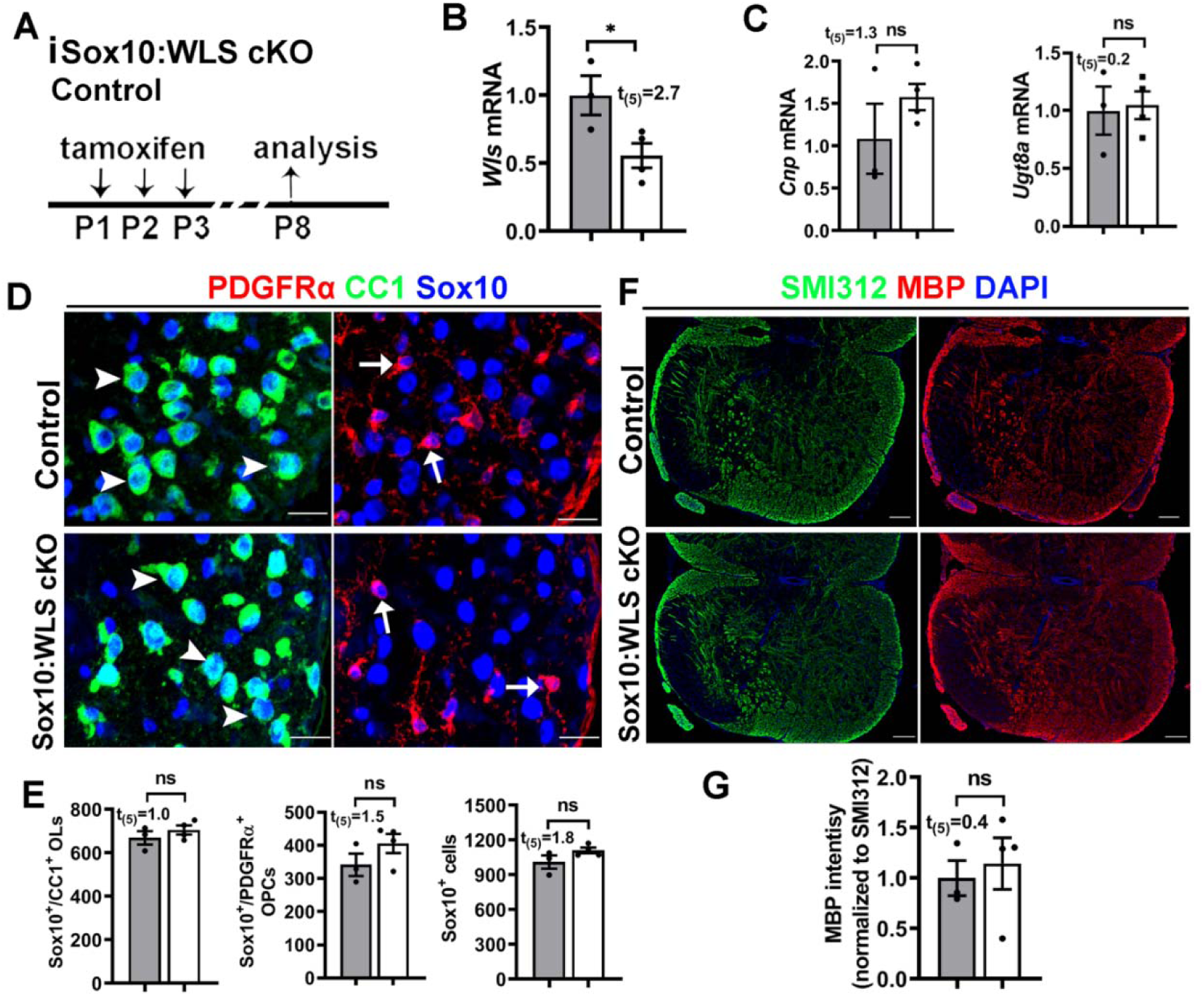
Disrupting WLS in Sox10-expressing oligodendroglial lineage cells does not perturb normal OPC differentiation and myelination. A, experimental design for **B-G**. Tamoxifen inducible *Sox10-CreER^T2^:Wls^m^* (iSox10:WLS cKO, n=4); littermate non-Cre control Wls^fl/fl^ mice (n=3). **B-C**, RT-qPCR quantification of Wls **(B)** and differentiated OL-enriched gene Cnp and Ugt8a **(C)** in the spinal cord. **D-E**, representative confocal images **(D)** and quantification **(E)** of Sox10^+^/CC1^+^ differentiated OLs (**D**, arrowheads), Sox10^+^/PDGFRα^+^ OPCs (**D**, arrows), and Sox10^+^ oligodendroglial lineage cells in the spinal cord. **F-G**, representative confocal images **(B)** and quantification **(C)** of myelination by MBP staining of the spinal cord. SMI312 signal was used as an internal control of MBP quantification. Scale bars, D, 20μm, E, 50μm.

### Sustained Sox9 activation mediates HIFα stabilization-elicited inhibition of OPC differentiation

The mRNA level of Sox9 was significantly upregulated in primary OPCs maintained in growth medium that had been treated with dimethyloxalylglycine (DMOG), a potent HIFα stabilizer (Yuen et al., 2014), and this upregulation was abrogated upon simultaneous treatment of chetomin (Fig. 11A), an inhibitor of HIFα transcriptional activity (Kung et al., 2004; Viziteu et al., 2016). To support the functional regulation, our assay of chromatin immunoprecipitation followed by qPCR assay (ChIP-qPCR) showed that HIFα physically bound to the promoter region (−828 bp) of the rat *Sox9* gene (Fig. 11B). We found that Sox9 mRNA expression was significantly upregulated in the spinal cord of PDGFRα:VHL cKO mice compared with littermate controls at P8 (Fig. 11C) and P14 (Fig. 11D). At the cellular level, sustained Sox9 expression was observed in HIFα-stabilized Sox10-expessing oligodendroglial lineage cells in Sox10:VHL cKO mice (Fig. 11E, right, arrowheads), in sharp contrast to control mice in which Sox9 was expressed at much lower level or barely detectable in Sox10-expressing cells (Fig. 11E, left, arrowheads). Our quantification showed that the percentage of Sox10^+^ cells expressing Sox9 (Fig. 11F) and the level of Sox9 mRNA (Fig. 11G) were significantly increased in Sox10:VHL cKO mice. Furthermore, Sox9 expression was significantly elevated in primary OPCs, which were purified from Sox10:VHL cKO neonatal brain (Fig. 11H). Sox9 is expressed in virtually all astrocytes in the postnatal CNS (Sun et al., 2017; Zhang et al., 2018b). We found that the density of Sox9^+^GFAP^+^ astrocytes was not significantly different in the spinal cord of Sox10:VHL cKO mutants from that in control mice (Fig. 11I). We also found that the percent of Sox10^+^ oligodendroglial cells expressing Sox9 (Fig. 11J1-J2) was unchanged in PDGFRα:HIFα cKO mice (Fig. 11J3).

**Figure 11.**
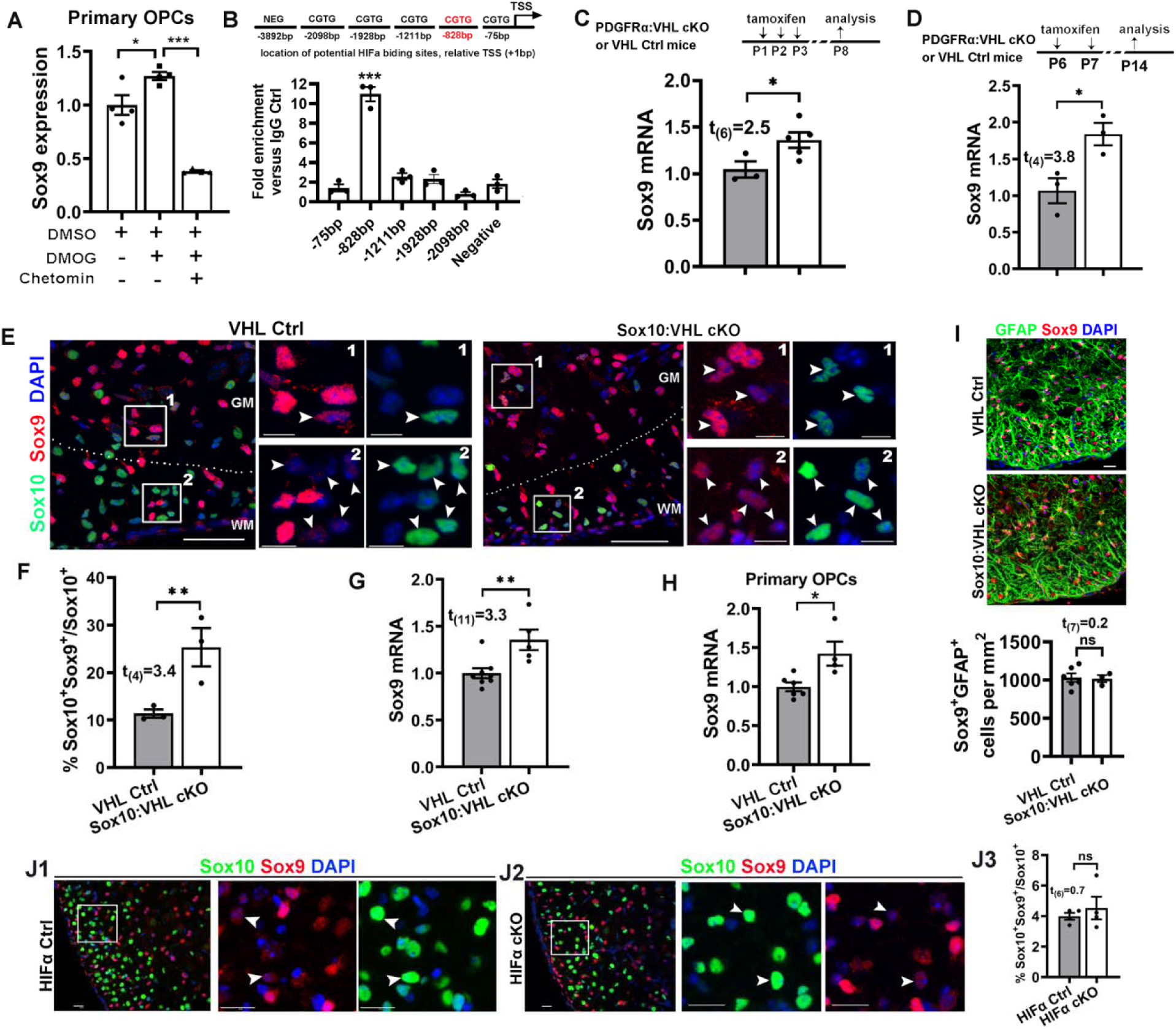
HIFα stabilization activates Sox9 expression. **A**, Sox9 expression in the purified primary OPCs assessed by RT-qPCR. Purified cortical OPCs from neonatal rat brains were treated with HIFα stabilizer dimethyloxalylglycine (DMOG, 1mM) in the absence and presence of HIFα signaling blocker chetomin (100nM) for 7 h. n=4 each group. One-way ANOVA followed by Tukey’s multiple comparisons, *F*_(2, 9)_ = 62.53 *P <* 0.0001. **B**, potential HIFα binding sites (CGTG) in the upstream sequence of rat Sox9 genes (^+^1bp is defined as the transcription start site, TSS) and ChIP-qPCR verification of physical binding of HIFα to the promoter region of -828bp in primary OPCs treated with DMOG. One way ANOVA with Dunnett post-hoc test comparing each group with negative control. *F*_(2,12)_ = 69.21 *P* < 0.0001. **C**, RT-qPCR quantification of Sox9 expression in P8 spinal cord of PDGFRα:VHL cKO (n=5) and littermate control(Vhl^fl/fl^, n=3) mice that had been treated with tamoxifen at P1, P2, and P3. **D**, RT-qPCR quantification of Sox9 expression in P14 spinal cord of PDGFRα:VHL cKO (n=3) and littermate control(Vh/^l/fl^, n=3) that had received tamoxifen at P6 and P7. **E**, confocal images showing that Sox9 is barely expressed in Sox10^+^ oligodendroglial lineage cells in the control spinal cord but upregulated in Sox10:VHL cKO mice at P5. WM, white matter, GM, gray matter. The boxed areas of #1 and #2 were shown at higher magnification at the right. **F**, percentage of Sox10^+^ oligodendroglial lineage cells expressing Sox9 in the spinal cord (n=3 each group). **G**, RT-qPCR quantification of Sox9 mRNA in the spinal cord of Sox10:VHL cKO (n=5) and littermate control (*Vhl*^fl/+^ and/or *Vhl*/^fl/fl^, n=8) mice at P5. **H**, RT-qPCR assay of Sox9 expression in primary OPCs isolated from the neonatal brain of *Sox10-Cre:Vhf^m^* (n=4) and littermate control (n=6) pups. **I**, representative confocal images and quantification of Sox9^+^GFAP^+^ astrocytes in the ventral spinal cord of P5 *Sox10-Cre:Vhl*^fl/fl^ (n=3) and littermate control (n=6) mice. J1-J3, representative confocal images and quantification of Sox10 and Sox9 in the ventral spinal cord of P8 PDGFRcrHIFα cKO (n=4) and littermate control (n=4) mice (tamoxifen at P1, P2, and P3). Arrowheads point to Sox10^+^Sox9^+^ cells. Scale bars, 5μm in E, 10 μm in I, 10 μm in boxed areas of panel E. 20 μm in J1-J2.

Recent data demonstrate that Sox9 is rapidly downregulated upon OPC differentiation (Reiprich et al., 2017), prompting us to hypothesize that sustained Sox9 expression may inhibits OPC differentiation. To test this hypothesis, we overexpressed Sox9 in primary OPCs in growth medium and assess OPC differentiation in differentiating medium (Fig. 12A). Sox9 mRNA and protein were significantly increased in primary OPCs transfected with Sox9-expressing plasmid compared with those transfected with empty vector (EV) (Fig. 12B-D). Sox9 overexpression reduced the mRNA expression of the most abundant myelin protein Plp and the potent pro­differentiation factor Myrf (Emery et al., 2009) assessed by RT-qPCR (Fig. 12 E), the number of ramified MBP^+^/Sox10^+^ oligodendrocytes assessed by immunocytochemistry (Fig. 12F), and the protein level of MBP assessed by Western blot (Fig. 12G). To define the role of Sox9 activation in HIFα-regulated OPC differentiation, we stabilized HIFα in OPCs by DMOG treatment and simultaneously knocked down Sox9 expression by siRNA (Fig. 12H-J). Our data showed that Sox9 downregulation rescued the degree of myelin gene inhibition elicited DMOG treatment, as evidenced by the density of MBP^+^/Sox10^+^ differentiated OLs with ramified morphology (Fig. 12K-L) and the expression of myelin gene *Mbp* (Fig. 12M). Collectively, our results suggest that sustained Sox9 expression is one of the downstream molecular pathways in mediating HIFα-regulated OPC differentiation.

**Figure 12.**
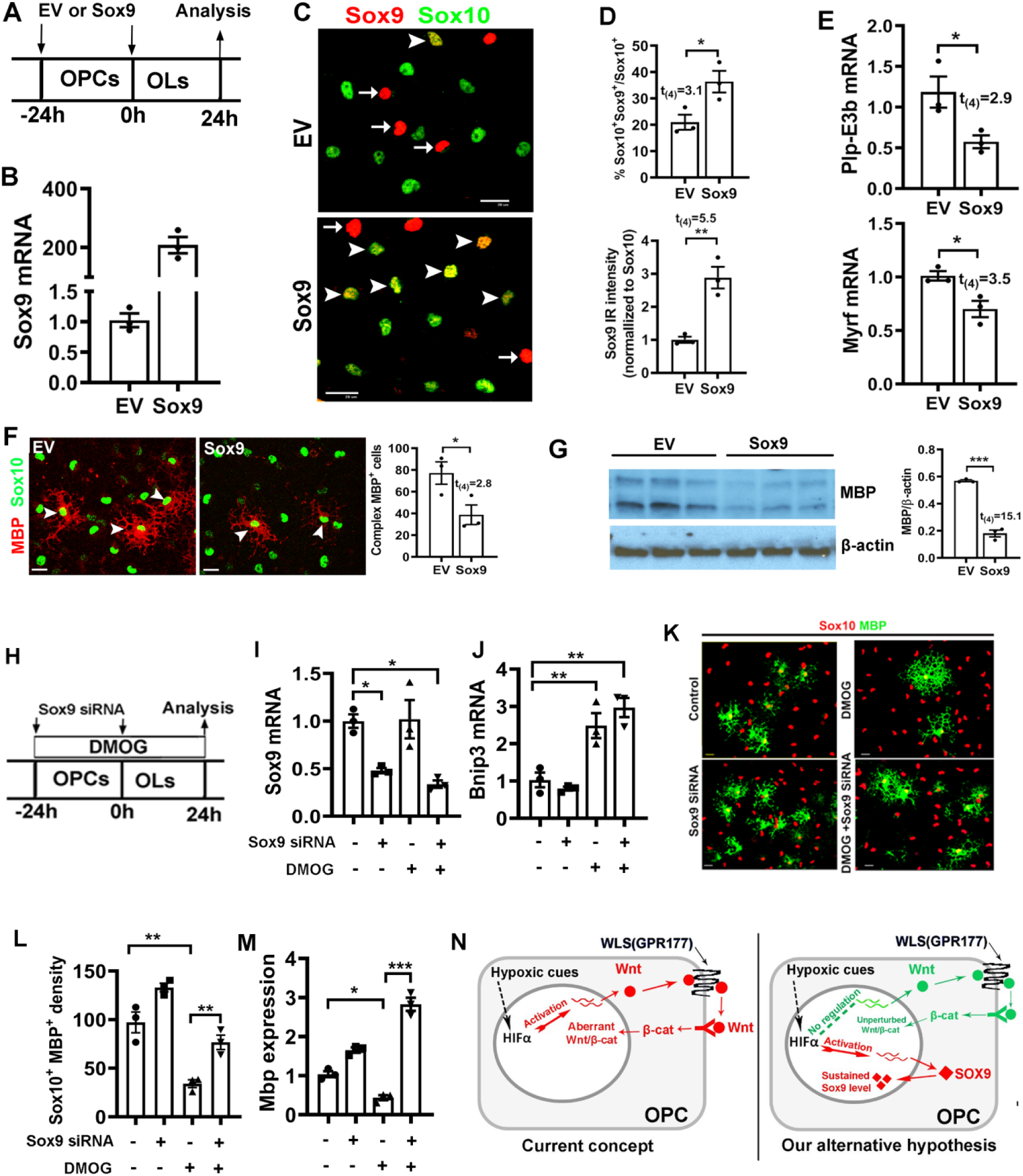
Sustained Sox9 expression inhibits OPC differentiation. **A**, experimental design for **B-G**. Cortical primary OPCs in growth medium were transfected with Sox9-expressing plasmid (Sox9, n=3) or empty vector (EV, n=3), then differentiate for 24 h in serum-free differentiating medium. **B**, RT-qPCR quantification of Sox9 mRNA expression. **C**, representative immunocytochemical images of Sox9/Sox10. Arrowheads point to Sox9^+^/Sox10^+^ oligodendrocytes and arrows point to Sox9^+^/Sox10^−^ cells, which are presumably astrocytes. Scale bar=2μm. **D**, percentage of Sox10^+^ oligodendroglial lineage cells that express Sox9 (top) and Sox9 immunoreactive intensity normalized to Sox10 (bottom). **E**, RT-qPCR assay of exon3b-containing *Plp (Plp-E3b)*, which specifically expresses in differentiated OLs, and myelin regulatory factor *(Myrf)*, a potent pro-differentiation gene. **F**, representative immunocytochemical images of MBPlSox10 and density (#/mm^2^) of Sox10^+^ oligodendroglial cells of complex ramified morphology indicated by MBP staining (arrowheads in **F**), Scale bar=1μm. **G**, Western blot images and quantification of MBP and internal loading control β-actin. **H**, experimental designs for panels **I-M**, primary OPCs were treated with HIFα stabilizer DMOG and transfected with Sox9 siRNA twice with 24 h apart. **I-J**, expression of *Sox9* **(I)** and HIFα target gene *Bnip3* **(J)** quantified by RT-qPCR. One way ANOVA followed by Tukey’s test; *F*_(3, 8)_ = 10.33, *P* = 0.004 *Sox9;* *F*_(3, 8)_ = 20.83, *P* = 0.0004 *Bnip3.* **K**, representative immunostaining images of differentiated OLs labeled by MBP and Sox10. Scale bar=10μm. **L-M**, density (#/mm^2^) of MBP^+^lSox10^+^ differentiated OLs **(L)** and RT-qPCR assay of *Mbp* mRNA expression **(M)**. One way ANOVA followed by Tukey’s test; *F*_(3, 8)_ = 33.29, *P* < 0.0001 MBP^+^/Sox10^+^; *F*_(3, 8)_ = 97.96, *P* < 0.0001 *Mbp.* **N**, previously proposed and our alternative working model explaining the role of HIFα in OPC differentiation. Endogenous HIFα regulates CNS myelination by controlling upstream OPC differentiation but not downstream OL maturation in perinatal and early postnatal CNS whereas aberrant HIFα activation (via VHL mutation of hypoxia-ischemia insult) inhibits OPC differentiation through the Wnt-independent and Sox9-dependent pathway.

## Discussion

### Summary of key findings and conclusions

Through the analysis of a series of HIFα genetic models (stage-dependent HIFα loss or gain-of-function in OPCs and OLs), our study provides compelling evidences supporting an alternative understanding of the cellular and molecular mechanisms underlying HIFα-regulated CNS myelination (Fig. 12N). At the cellular level, our in vivo genetic data support a new working model in which physiological HIFα transiently regulates OPC differentiation whereas persistent HIFα activation directly causes arrested OPC differentiation, leading to developmental hypomyelination. At the molecular level, HIFα regulates OPC differentiation in a manner independent of the autocrine Wnt/β-catenin signaling. Instead, dysregulated Sox9 expression is one of the downstream pathways in mediating HIFα activation-elicited inhibition of OPC differentiation (Fig. 12N).

### HIFα regulates developmental myelination by controlling OPC differentiation but not subsequent oligodendrocyte maturation

One of the unexpected observations is the differential responses of OPCs and OLs to HIFα disruption and hyperactivation. Disrupting or stabilizing HIFα function results in severe disturbance of myelin gene expression and developmental myelination in the CNS of OPC-specific PDGFRα:HIFα (VHL) cKO mice. In sharp contrast, HIFα disruption or stabilization in Cnp:HIFα (VHL) cKO mice does not perturb normal myelin gene expression and myelination.

Previous genetic studies suggest that *Cnp-Cre* elicits gene disruption primarily in the later stages of oligodendrocyte development. For example, disrupting DNA methyltransferase 1 (DNMT1) by *Cnp-Cre* line shows no phenotypes in OPC proliferation and expansion, in sharp contrast to DMNT1 disruption by *Olig1-Cre* which targets the progenitor stages of the oligodendroglial lineage (Moyon et al., 2016). Our recent studies also demonstrated that disrupting Sox2 by *Cnp-Cre* shows no phenotypes in OPC proliferation and expansion, in sharp contrast to Sox2 disruption by *Sox10-Cre* or *Pdgfrα-CreER^T2^* which targets OPCs (Zhang et al., 2018a; Zhang et al., 2018b). Therefore, the discrepant phenotypes in HIFα (or VHL) deletion driven by Cnp-Cre versus *Sox10-Cre* or *Pdgfrα-CreER^T2^* (Fig. 2, 4, 5) support the interpretation that HIFα play a major role in OPC differentiation into OLs and is dispensable for oligodendrocyte maturation.

We observed impaired OPC differentiation in HIFα activation (Fig. 2A-C, Fig. 3) and inactivation (Fig. 4A-D) genetic models. These observations suggest delicate downstream pathways regulated by HIFα. We recently reported that HIFα depletion reduces the canonical HIFα target genes involving in glucose metabolism and CNS angiogenesis (Zhang et al., 2020), both of which entails energy supply for OPC differentiation(Yuen et al., 2014). It is possible that HIFα inactivation may transiently affect OPC differentiation through limiting the energy supply regulated by physiological HIFα whereas sustained HIFα activation might stimulate additional non-canonical HIFα target genes which are otherwise non-responsive to physiological HIFα during normal development. Our experimental data identified one of such non-canonical targets as Sox9 whose expression is upregulated in HIFα activation mice and unaltered in HIFα inactivation mice. Furthermore, the unaltered developmental myelination in OL-specific HIFα activation and inactivation models indicates that OLs are resistant to HIFα dysregulation, which is in line with the previous concept that OLs are remarkably resistant to hypoxia/ischemia injury (Back et al., 2002; Back et al., 2007).

### Molecular mechanisms underlying HIFα-regulated OPC differentiation

Previous study employed cerebellar slice/cell culture as a studying model and proposed that the autocrine activation of Wnt/β-catenin signaling in OPCs by Wnt7a/Wnt7b plays a crucial role in HIFα activation-induced hypomyelination (Yuen et al., 2014). Our data derived from *in vivo* genetic models do not support this concept. Firstly, unbiased RNA-seq analysis did not reveal dysregulation of the Wnt/β-catenin signaling activity and Wnt7a/Wnt7b expression. Secondly, neither the activity of Wnt/β-catenin signaling nor the expression of Wnt7a/Wnt7b is perturbed in HIFα-stabilized OPCs. Thirdly, disrupting the Wnt secretion mediator WLS blocks OPC-derived autocrine Wnt/β-catenin signaling, but it does not affect the degree of inhibited OPC differentiation elicited by HIFα hyperactivation. Our findings indicate that Wnt7a/Wnt7b or Wntl3-catenin signaling is unlikely a direct target of HIFα in the oligodendroglial lineage cells (Fig. 12N), which is supported by a recently submitted study from an independent group (Allan et al., 2020). The reasons for the different observations between our current study and prior study (Yuen et al., 2014) are unclear. It is possible that non-cellular-specificity and/or off-target of pharmacological compounds may underlie the discrepancy. HIFα stabilization induced by hypoxia treatment or by pharmacological DMOG application occurs not only in oligodendroglial lineage cells but also in other lineages of neural cells or vascular cells. Similarly, Wntl3-catenin inhibition in cultured slices by pharmacological XAV939 or IWP2 treatment may also happen in other lineage cells (Yuen et al., 2014). We used cell-specific *Cre-loxP* genetic approaches to circumvent the potential caveats of pharmacological application and convincingly demonstrated that OPC-derived autocrine Wnt/β-catenin plays a minor, if any, role in mediating HIFα-regulated OPC differentiation and myelination.

Another novel finding of this study is the identification of Sox9 as a potential target that mediates HIFα stabilization-elicited inhibition of OPC differentiation (Fig. 12N, right panel). Sox9 is highly expressed in neural stem cells (NSCs) (Scott et al., 2010), downregulated in OPCs (Stolt et al., 2003), and completely absent from differentiated OLs (Sun et al., 2017). Sox9 disruption affects OPC specification but not OPC terminal differentiation (Finzsch et al., 2008; Stolt et al., 2003). HIFα stabilization activated Sox9 expression in OPCs both *in vitro* and *in vivo*, which is in agreement with previous data showing that Sox9 is transcriptionally activated by HIFα in chondrocytes (Amarilio et al., 2007) and pancreatic β-cell precursor cells (Puri et al., 2013). We found that elevated Sox9 expression perturbed OPC differentiation and that Sox9 knockdown rescued the extent of inhibited OPC differentiation elicited by pharmacological HIFα stabilization in primary OPC culture system.

It should be pointed out that the non-regulation of HIFα-Wnt axis does not negate the crucial role of Wnt/β-catenin in oligodendroglial lineage cells. Previous studies have revealed an inhibitory effect of Wnt/β-catenin activation on OPC differentiation (Guo et al., 2015). However, these studies modulated the signaling activity through manipulating the intracellular components or relevant regulatory molecules within OPCs, for instance, by disrupting intracellular β-catenin (Fancy et al., 2009; Feigenson et al., 2009; Ye et al., 2009), Axin2 (Fancy et al., 2011), APC (Fancy et al., 2014; Lang et al., 2013), APCDD1 (Lee et al., 2015b), and Daam2 (Lee et al., 2015a). The Wnt-producing cells, which play a crucial role in activating the intracellular Wnt/β-catenin signaling axis in OPCs remain unknown (Guo et al., 2015). Our WLS cKO data demonstrate that oligodendroglial lineage-derived autocrine Wnt plays a minor role in OPC differentiation and myelination in the early postnatal CNS under physiological (Fig. 10) or HIFα-stabilized (Figs. 8, 9) conditions, suggesting that Wnt ligands derived from other lineage cells may be responsible for control OPC differentiation through paracrine Wnt signaling. Previous RNA-seq data (Zhang et al., 2014) indicate that astrocytes may be the major cellular source of Wnt proteins, for example Wnt7a and Wnt5a. Future studies are needed to test the tempting hypothesis that astrocyte-derived paracrine Wnt signaling may play major a role in controlling OPC differentiation under normal or HIFα-stabilized conditions.

### Implications of our findings in hypoxia-ischemia-induced oligodendroglial pathology in preterm white matter injury (WMI)

Disturbance of normal developmental myelination is one of the established pathological hallmarks in preterm (or premature) infants affected by diffuse WMI (Back, 2017; Deng, 2010). Hypoxia/ischemia-induced diffuse WMI frequently occurs in the brain of preterm infants because of the anatomical and physiological immaturity of the respiratory system and the brain white matter vasculature. We demonstrated that, compared with healthy controls, HIF1α was markedly elevated in the brain white matter of mice subjected to hypoxia-ischemia injury at the time equivalent to developmental myelination in human brains (Fig. 1F-I). Our findings -sustained HIFα activation disturbs OPC differentiation and results in hypomyelination -suggest that oligodendroglial HIFα upregulation may play a crucial role in hypoxia-ischemia-induced myelination disturbance. The function and molecular mechanisms of oligodendroglial HIFα in hypoxia-ischemia-induced hypomyelination (Liu et al., 2011) have yet to be determined in genetic animal models. Our *in vivo* data indicate that sustained HIFα activation in response to hypoxia-ischemia injury may inhibit OPC differentiation and cause hypomyelination and that the underlying mechanisms may be independent of OPC autocrine canonical Wnt signaling as previously proposed. Future studies using cell-specific genetic HIFα models (HIFα loss and gain-of-function) and pre-term equivalent mouse models of diffuse white matter injury are needed to address these important questions.

## Competing interests

The authors declare no competing financial interests

## Acknowledgements

This study was supported by the grants funded by NIH/NINDS (R21NS109790 and R01NS094559 to F.G.) and Shriners Hospitals for Children (86100, 85200-NCA16, 85107-NCA-19 to F.G., 84307-NCAL to S.Z.).

## Notes

### Competing Interest Statement

The authors have declared no competing interest.

### Summary of Updates

We reorganize section sequences and added more data in our revised paper. Fig. 1-5 revised. author and author affiliation updated; disscusion section updated.

